# Mechanisms of phenotypic trade-offs in the resource acquisition-allocation Y-model

**DOI:** 10.1101/2025.07.07.662185

**Authors:** Krish Sanghvi, Samuel J. L. Gascoigne, Irem Sepil

## Abstract

Phenotypic trade-offs, predicted to occur due to resource constraints, are not commonly observed. This discrepancy is often explained by the Y-model, which shows that greater variation in resource acquisition (CV_*A*_) than variation in allocation strategy (CV_*X*_) masks among-individual phenotypic trade-offs. However, the Y-model is a heuristic rather than a quantitative tool with testable predictions. Additionally, the mechanisms modulating trade-offs in the model remain unclear. Here, we simulate different parameters of the Y-model to understand their influence. We find that ‘CV_*A*_/(CV_*A*_ + CV_*X*_)’ accurately predicts the direction and strength of phenotypic correlations. Contrary to common assumptions, the mean resource acquired by a population does not influence trade-offs; instead, the mean allocation strategy of a population does. Importantly, within-individual dependence of allocation on acquisition exacerbates the influence of mean allocation. This causes greater sensitivity of phenotypic correlations to changes in CV_*A*_ or CV_*X*_, compared to independent resource acquisition and allocation strategies. We generate novel, testable hypotheses about trade-offs in the context of dietary restriction, plasticity, polymorphism, and pace-of-life, and validate our model with empirical data to demonstrate its robustness. By systematically partitioning the influence of means, variances, and covariance, in the acquisition-allocation Y-model, we provide a simple, generalisable, quantitative synthesis for understanding phenotypic trade-offs.

## Introduction

Organisms have evolved to acquire resources from the environment and allocate these toward different life-history traits such as growth, survival, and reproduction, which collectively improve fitness (Lailvaux and Husak, 2014). Individuals often vary in acquired resources and their strategies to partition these towards different traits (Healy et al, 2019; Partridge and Sibly, 1991; Salzman et al, 2018). For example, ‘slow-lived’ organisms allocate relatively more energy towards improving late-life survival while fast-lived ones allocate more toward early-life reproduction (Healy et al, 2019; Montiglio et al, 2018). Because resources available to organisms are finite, a fundamental prediction in life-history theory is that trade-offs exist between the fraction of acquired resources allocated to each trait, thereby creating negative phenotypic correlations between life-history traits (reviewed in Agrawal, 2019; Gascoigne et al, 2023; Maklakov and Chapman, 2019; Ricklefs and Wikelski, 2002; Stearns, 1989, 1992).

Phenotypic trade-offs among life-history traits have been reported across taxa (reviewed in Edward and Chapman, 2011; Reznick, 1985; Roff, 1996; Zera and Harshman, 2001), through the quantification of phenotypic and genetic correlations (e.g. Attisano et al, 2012; Roff, 2000; Roff et al, 2007; Srgo and Hoffman, 2004), the experimental manipulation of individual traits (e.g. Gascoigne et al, 2022; Moczek and Nijhout, 2004), and via selection experiments. For instance, individuals that produce more offspring earlier in life, are found to live shorter lives and produce relatively fewer offspring at old ages (Descamps et al, 2006; Flatt, 2011; Hayward et al, 2015; Lemaitre et al, 2015; Reznick, 1983; Sanghvi et al, 2022; Young et al, 2024). Furthermore, experimental removal of the germline improves the ability of individuals to repair somatic damage (Chen et al, 2020) and extends their longevity (Cox et al, 2010), compared to those with intact germlines. This represents trade-offs between reproduction and somatic maintenance. Experimental evolution studies also show that individuals in populations selected to live longer, also reproduce later (Gasser et al, 2000; Leroi et al, 1994; Miyatake, 1997) or reproduce less (e.g. Maklakov et al, 2017), compared to those in populations selected to live shorter lives. Yet, despite some evidence for phenotypic trade-offs and predictions of their ubiquity, more often than not, studies do not observe trade-offs (e.g. Robert et al, 2015; Sanghvi et al, 2025a; reviewed in Bolund et al, 2020; Lim et al, 2014; Reznick et al, 2000). In fact, recent meta-analyses show that life-history traits co-vary positively among-individuals at a phenotypic (Have-Audet et al, 2022; Winder et al, 2025) and genotypic level (Chang et al, 2024) across the animal kingdom.

Van Noordwijk and De Jong (1986) formally established the fundamental theoretical framework used for understanding phenotypic trade-offs, in their ‘Y-model’. This model conceptualized resource acquisition and its allocation between two traits, providing a framework for when among-individual phenotypic trade-offs might be observed or masked (Figure S1). Crucially, it explained why life-history traits often exhibit positive phenotypic correlations despite underlying trade-offs in resource partitioning within an individual (also see Saglam et al 2008; De Jong and Van Noordwijk 1992; Laitinen and Nikoloski 2024; Robinson and Beckman 2013). The model’s core prediction states that greater variation in allocation increases the detectability of phenotypic trade-offs, while greater variation in acquisition buffers trade-offs (De Jong 1993; De Laguerie et al 1991; Descamps et al 2016; Hashemi et al 2024; Haave-Audet et al 2022; Johnson and Nasrullah 2024; Metcalf 2016; Reznick et al 2000; Roff and Fairbairn 2007; Schluter et al 1991; Zera and Harshman 2001). While this prediction has been tested by some studies (e.g. Boggs 2009; Boggs and Freeman 2005; Brown 2003; Christians, 2000; Douhard et al 2021; Glazier 1999; King et al 2011; Messina and Fry 2003; Robinson and Beckerman 2013; Smiseth et al 2014; Tatar and Carey 1995), it does not universally hold (reviewed in Roff and Fairbairn 2007).

There are various reasons for discrepancies between predictions of the Y-model versus empirical results. First, these discrepancies occur due to the difficulty in measuring acquisition and allocation, and quantifying them in the same units (Roff and Fairbairn, 2007). Second, simplistic assumptions of the Y-model make it biologically unrealistic (Cohen et al, 2020; Johnson and Nasrullah 2024), hence only a handful of theoretical studies (Descamps et al, 2016; Roff and Fairbairn, 2007) have used empirical data to validate their models. Third, discrepancies occur because the Y-model only explains *why* phenotypic trade-offs might be detected or masked, but cannot yet predict the strength of the phenotypic correlation. This prevents empirical studies from generating quantitative hypotheses *a priori*, limiting the model’s utility to being a heuristic that’s explains empirical results post-hoc (e.g. Forrester et al, 2025; Sanghvi et al, 2025a). Fourth, studies often misinterpret the Y-model (reviewed in Laskowski et al, 2021; Roff and Fairbairn, 2007) by conflating the effects of its different parameters, namely: variation in resource acquisition and allocation, mean resource acquisition and allocation (e.g. Attwood, 2025; Douhard et al, 2021; Smiseth et al, 2014), and their covariance. The independent effects of some of these parameters on phenotypic trade-offs have been previously clarified, for example, the influence of acquisition-allocation covariance (Descamps et al, 2016; Robinson and Beckerman, 2013; Zajitschek and Connallon, 2017), or of variation in acquisition or allocation (Johnson and Nasrullah 2024; Roff and Fairbairn, 2007; Van Noordwijk and De Jong, 1986). However, misinterpretation persists because the direct, independent influence of mean acquisition and allocation remains unclear. Additionally, how mean acquisition (i.e. average population-level resource richness), mean allocation (i.e. average population-level life-history strategy to partition resources), their covariance (i.e. dependence of allocation on acquisition), and their variance (between-individual differences), interact to influence among-individual phenotypic correlations in a population, remains unanswered.

Here, we use simulations to extend the seminal ‘Y-model’ (Van Noordwijk & De Jong, 1986) to better understand the mechanisms that create phenotypic trade-offs. We address three specific aims: (aim 1) Quantify how relative variation in resource allocation and acquisition impact among-individual phenotypic correlations (Figure S2); (aim 2) Test the direct impact of mean allocation strategy of, and mean resources acquired by, a population, on phenotypic correlations (Figure S3); (aim 3) Investigate whether the covariance between acquisition and allocation modulates phenotypic correlations (Figure S4). Through these aims, we systematically separate the independent and interactive influence of each Y-model parameter on phenotypic trade-offs; validate our models against empirical data; generate novel quantitative hypotheses about trade-offs; discuss how these can be empirically tested by incorporating biological realism; and provide solutions for interpreting the Y-model.

## Method

### Simulation overview

We used agent-based simulations to understand how resources acquired and the proportion of these allocated towards two traits, impact among-individual phenotypic correlations between these two traits. All analyses were conducted on R.v.4.3 (R core team, 2020). We simulated a simple organism that allocated its entire common pool of acquired resources towards only two traits, and assumed that the amount of resources allocated to a trait corresponded to the phenotypic measurements that trait. We modelled these as:

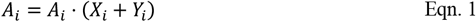

thus,

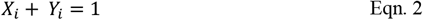

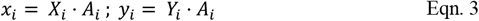

*A*_*i*_ Eqn. 3 where, *A*_*i*_ was the total amount of resource acquired, *X*_*i*_ was the proportion of that resource allocated to trait *x*, and *Y*_*i*_ was the proportion allocated to trait *y*, by the *i*^*th*^ individual. For the *i*^*th*^ individual, *x*_*i*_ and *y*_*i*_ represented the absolute units of resources invested in traits *x* and *y*, thus their phenotypic measurements, respectively. To create variation between-individuals in acquisition and allocation, we simulated coefficients of variation (CV) of resource acquisition (CV_*A*_) and of the proportion of resources allocated to trait *x* (CV_*X*_) as:

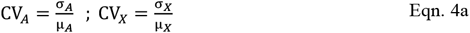

thus,

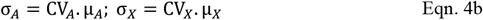

where μ_*A*_ and σ_*A*_ were the sample distribution mean and standard deviation for resource acquisition, while μ_*X*_ and σ_*X*_ were the sample distribution mean and standard deviation for the proportion of resources allocated to trait *x*, respectively. In models 1 and 2, the resources acquired and allocated by the *i*^*th*^ individual were sampled from independent normal distributions (Ɲ) using a Monte-Carlo process (Figure S5, S6, S7) as:

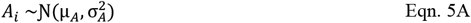

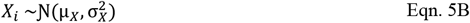

For all models, *A*_*i*_ was bounded by and truncated at zero, while *X*_*i*_ was between zero and one (Figure S5, S6). For every *i*^*th*^ individual, after sampling the proportion of resources allocated to trait *x* (*X*_*i*_), we sampled the proportion allocated to trait *y* as:

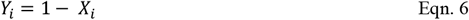

### Aim 1

In aim 1 (Model 1), we tested how variation in acquisition and allocation influence phenotypic correlations. For this, we fixed the values of μ_*A*_ and μ_*X*_, while varying σ_*A*_ and σ_*X*_ to create different values of CV_*A*_ and CV_*X*_; *A*_*i*_ and *X*_*i*_ did not co-vary (Figure S7). Due to the inverse relationship between *X* and *Y*, within-individuals, the proportion of resources allocated to each trait always correlated negatively (i.e. Cor(*X,Y*) = -1). We simulated a population of 2500 individuals for every unique combination of CV_*A*_ and CV_*X*_ values, where each CV was iterated between 0.005 to 0.25, in intervals of 0.005 (Table 1). This range of simulated CV_*A*_ and CV_*X*_ represents biologically meaningful variation given the boundedness and type of data (Acasuso-Rivero et al, 2019; Araya-Ajoy et al, 2023; Botta-Dukat, 2023; Garcia-Gonzalez et al, 2012; Guadard et al, 2019). The value of μ_*A*_ was fixed as 100; μ_*X*_ was fixed at 0.5 implying 50% of resources were allocated to each trait when averaged across the population (Table 1). When *X* + *Y* = 1, σ_*X*_ = σ_*Y*_ and 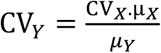. Therefore when μ _*X*_ = μ_*Y*_ = 0.5, CV_*X*_ = CV_*Y*_ (Figure S8, S9). For each unique pairwise combination of CV_*A*_ and CV_*X*_, we retrospectively calculated the among-individual phenotypic correlation between traits *x* and *y* as:

**Table 1:** Summary of parameters values in models 1-4. Values for mean (μ) and coefficient of variation (CV) of resource acquisition (*A*) and allocation strategy (*X*), as well as their correlation, Cor(*A,X*).

| Model ID | $CV_A$ | $CV_X$ | $\mu_A$ | $\mu_X$ | $\text{Cor}(A, X)$ |
| --- | --- | --- | --- | --- | --- |
| 1 | 0.005<br>to 0.25 | 0.005<br>to 0.25 | 100 | 0.5 | 0 |
| 2A | 0.005<br>to 0.25 | 0.005<br>to 0.25 | 1, 10, 100, 1000, 10000 | 0.5 | 0 |
| 2B | 0.005<br>to 0.25 | 0.005<br>to 0.25 | 100 | 0.1, 0.4, 0.7 | 0 |
| 3A | 0.005<br>to 0.25 | 0.005<br>to 0.25 | 1, 10, 100, 1000, 10000 | 0.5 | 1 |
| 3B | 0.005<br>to 0.25 | 0.005<br>to 0.25 | 100 | 0.1, 0.4, 0.7 | 1 |
| 4 | 0.005<br>to 0.025 | 0.005<br>to 0.025 | 1, 100, 10000 | 0.1, 0.4, 0.5, 0.7 | 0, 1, -1 |

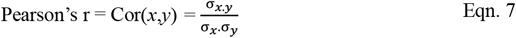

This correlation quantified the strength of the linear association between absolute amounts of resources invested in traits *x* and *y*, i.e. their phenotypic measurements *x*_*i*_ = *X*_*i*_ · *A*_*i*_ and *y*_*i*_ = *Y*_*i*_ · *A*_*i*_, respectively, across all 2500 individuals in a population. Some examples of among-individual phenotypic correlations for different CV_*A*_ and CV_*X*_ values are shown in Figure S10.

### Aim 2

We investigated whether the average amount of resources acquired by a population (μ_*A*_), impacts phenotypic correlations between traits *x* and *y*, in addition to the influence of CV_*A*_ and CV_*X*_ (Model 2A). For this, we kept the simulation structure identical to model 1 where we sampled *A*_*i*_ and *X*_*i*_ from various distributions, each with different combinations of values for CV_*A*_ and CV_*X*_. However, in model 2A, we incorporated one change. Specifically, we iterated five different values of μ_*A*_: 1, 10, 100, 1000, and 10000 (Figure S11), but kept μ_*X*_ fixed at 0.5 (Table 1). For each unique combination of CV_*A*_, CV_*X*_, and μ_*A*_, we simulated a population of 2500 individuals. Comparing results of model 2A to 1 would thus allow us to isolate the effects of μ_*A*_ from the influence of variation in allocation and acquisition, on the among-individual phenotypic correlation between *x* and *y*.

Next, we tested whether the mean resource partitioning strategy of a population, μ_*X*_, impacts the phenotypic correlation between traits *x* and *y* in addition to the influence of CV_*A*_ and CV_*X*_ (Model 2B). For this, we iterated three different values of μ_*X*_: 0.1, 0.4, 0.7, (with corresponding μ_*Y*_ values being 0.9, 0.6, and 0.3 respectively; Figure S12A, S12B), but kept the rest of the simulation structure and parameters identical to model 1. Specifically, μ_*A*_ was fixed at 100, CV_*A*_ and CV_*X*_ were iterated between 0.005 and 0.25 (Table 1). For each unique combination of CV_*X*_, CV_*A*_, and μ_*X*_, we simulated 2500 individuals. Comparing results of model 2B to model 1 thus allowed us to isolate the effects of μ_*X*_ on phenotypic correlations between *x* and *y*. In 2B, because μ_*X*_ ≠ μ_*Y*_, therefore, CV_*X*_ ≠ CV_*Y*_ (Figure S13). We ensured that in models 1, 2A and 2B, there was no covariance between allocation and acquisition (Figure S14).

### Aim 3

To understand whether within-individual dependence of allocation strategy (*X*_*i*_) on resources acquired (*A*_*i*_) impacts the phenotypic correlation between traits *x* and *y*, we sampled *A*_*i*_ and *X*_*i*_ from a bivariate normal distribution of the form:

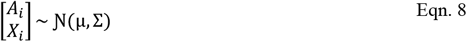

with a mean of 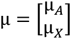 and a variance-co-variance matrix given by:

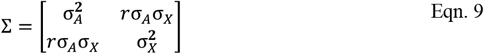

This allowed creating correlated values for *A*_*i*_ and *X*_*i*_. Using this matrix, we created two models. In model 3A, we specified a positive correlation (*r* = 1) between acquisition and allocation, i.e. Cor(*A,X*) *=* 1 (Figure S15), while iterating mean resource acquisition (μ_*A*_). μ_*A*_ got the values 1, 10, 100, 1000, 10000, while μ_*X*_ was held constant at 0.5. In model 3B, we specified *r* = 1 again, but instead we iterated μ_*X*_ (0.1, 0.4, and 0.7) while holding μ_*A*_ constant at 100 (Figure S16, S17; Table 1). Alike models 1, 2A, and 2B, in models 3A and 3B, we simulated different unique combinations of CV_*A*_ and CV_*X*_ values each ranging from 0.005 and 0.25 in intervals of 0.005, in order to sample allocation and acquisition. Similarly, in 3A and B, we simulated a population of 2500 individuals for every unique combination of CV_*X*_, CV_*A*_, μ_*X*_ and μ_*A*_. Comparing results of model 3A and 3B to models 1, 2A, and 2B allowed isolating effects of Cor(*A,X*) from effects of CV_*X*_, CV_*A*_, μ_*A*_, and μ_*X*_,. Negative correlations between allocation and acquisition had identical meaning to positive correlations between them (Figure S18) except that here, the identity of the focal allocation trait would be swapped (see Supplementary section S1; Figure S19; model 4). Therefore, model 3 could be interpreted as having either positive or negative covariance between acquisition and allocation because when Cor(*A,X*) = -1, Cor(*A,Y*) = 1 and vice versa.

### Model diagnostics

For models 1-3, we ran diagnostics to ensure that the specified CV and means of, and covariance between, acquisition and allocation, accurately matched the simulated parameter values (Figures S5-S17; Table 1).

### Statistically quantifying effects

We constructed a fourth simulation model (Model 4) where we simultaneously varied the values of CV_*A*_, CV_*X*_, μ_*A*_, μ_*X*_, and Cor(*A,X*). We parameterised these as follows: CV_*X*_, CV_*A*_ ranged from 0.005 to 0.25 in steps of 0.005; μ_*A*_ was 1, 100 or 10000; μ_*X*_ got the values 0.1, 0.4, 0.5, or 0.7; and Cor(*A,X*) was 0, -1, or 1 (Table 1). For each unique combination of CV_*A*_, CV_*X*_, μ_*A*_, μ_*X*_, and Cor(*A,X*) values, we simulated a population of 2500 individuals, and then retrospectively calculated the phenotypic correlation between traits *x* and *y*. While models 1-3 isolated the effects of each parameter from others, model 4 allowed statistical quantification of the interactive effects of the parameters on phenotypic correlations through linear models (*lme4* package, Bates et al, 2014). In separate linear models, we specified various combinations of additive and interactive effects of the parameters as fixed effects. Here, the among-individual phenotypic correlation, Cor(*x,y*), calculated separately for each population of 2500 individuals with a unique set of parameter values, was the dependent variable (Table 2). Then, using Akaike Information Criteria (AIC), we selected the best fitting linear model to understand the statistical significance of each parameter on Cor(*x,y*). This model covered 90000 unique points in the parameter space.

**Table 2:** Linear model terms to test how different parameters influence phenotypic correlations in model 4. Values of all parameters: μ_*X*_, μ_*A*_, CV_*X*_, CV_*A*_, and Cor(*A,X*), are varied. Linear model with lowest AIC was chosen as the best fitting (grey). R^2^ quantifies the % variation in Cor(*x,y*) explained by the linear model terms. ‘:’ implies interaction.

| Linear model terms | AIC | Model $R^2$ | DF |
| --- | --- | --- | --- |
| $(CV_A/CV_X) + \mu_A + \mu_X + \text{Cor}(A, X)$ | 193438 | 26.5% | 6 |
| $(CV_A:CV_X) + \mu_A + \mu_X + \text{Cor}(A, X)$ | 118792 | 67.9% | 8 |
| $[CV_A/(CV_A + CV_X)] + \mu_X + \mu_A + \text{Cor}(A, X)$ | 107523 | 71.7% | 6 |
| $[CV_A/(CV_A + CV_X)] + \mu_A + [\mu_X: \text{Cor}(A, X)]$ | 84308 | 78.2% | 7 |
| $[CV_A/(CV_A + CV_X)] + [CV_A/(CV_A + CV_Y)] + \mu_A + \mu_X: \text{Cor}(A, X)$ | 72729 | 80.8% | 8 |
| $[CV_A/(CV_A + CV_X)] + [CV_A/(CV_A + CV_Y)] + [\mu_X: \text{Cor}(A, X)]$ | 72727 | 80.8% | 7 |
| $[CV_A/(CV_A + CV_X)]: \mu_X: [\text{Cor}(A, X)] + [CV_A/(CV_A + CV_Y)]$ | 64059 | 82.6% | 10 |

### Proof of concept using complex distributions

To ensure that our results were not specific to sampling from normal distributions, we also created simulations where acquisition was sampled from a Gamma distribution and allocation from a Beta distribution. Results from these models agreed with results from normal distribution models (Supplementary section S2, Figure S20, S21). While these distributions are more biologically realistic, their parameters are harder to interpret. Additionally, these complex distributions did not generate distinct predictions compared to models 1-4. Therefore, only normal distributions were implemented across our simulations.

### Validation with empirical data

To demonstrate the utility of our simulations, we obtained empirical data on the parameters CV_*A*_, CV_*X*_, μ_*A*_, μ_*X*_, and Cor(*A,X*) from Table 1 from King et al (2011b). This classic study’s data has been used for over a decade by theoreticians to test Y-model predictions (in King et al, 2011b, Robinson and Beckerman, 2013). Using these data, we simulated a population of individuals (Supplementary section S3), and then predicted the among-individual phenotypic correlation. We then compared these simulated phenotypic correlations predicted by modelling the five parameters, to their empirically observed phenotypic correlations (Supplementary section S3). Furthermore, to test the influence of each parameter on the phenotypic correlation, we independently changed the value of one parameter at a time in different iterations of the model. We then compared the simulated phenotypic correlation generated using altered parameters, to the simulated phenotypic correlation generated with original parameter values, and to the empirical correlation (Supplementary section S3).

## Results

### Aim 1

In model 1, where only coefficients of variation of acquisition and allocation were iterated, we found that irrespective of their values, when CV_*A*_ > CV_*X*_, traits *x* and *y* correlated positively; when CV_*A*_ < CV_*X*_, traits *x* and *y* correlated negatively (Figure 1A); and when CV_*A*_ = CV_*X*_, the phenotypic correlation was zero (Figure S10). Interestingly, the phenotypic correlation, Cor(*x,y*), could be expressed as a function of ‘CV_*A*_/(CV_*A*_ + CV_*X*_)’ (Figure 1B). While Cor(*x,y*) was also a function of CV_*A*_/CV_*A*_, interpreting this was less intuitive due to asymmetry and extreme changes in the function’s shape (Figure 1C, further elaborated in model 4).

**Figure 1:**
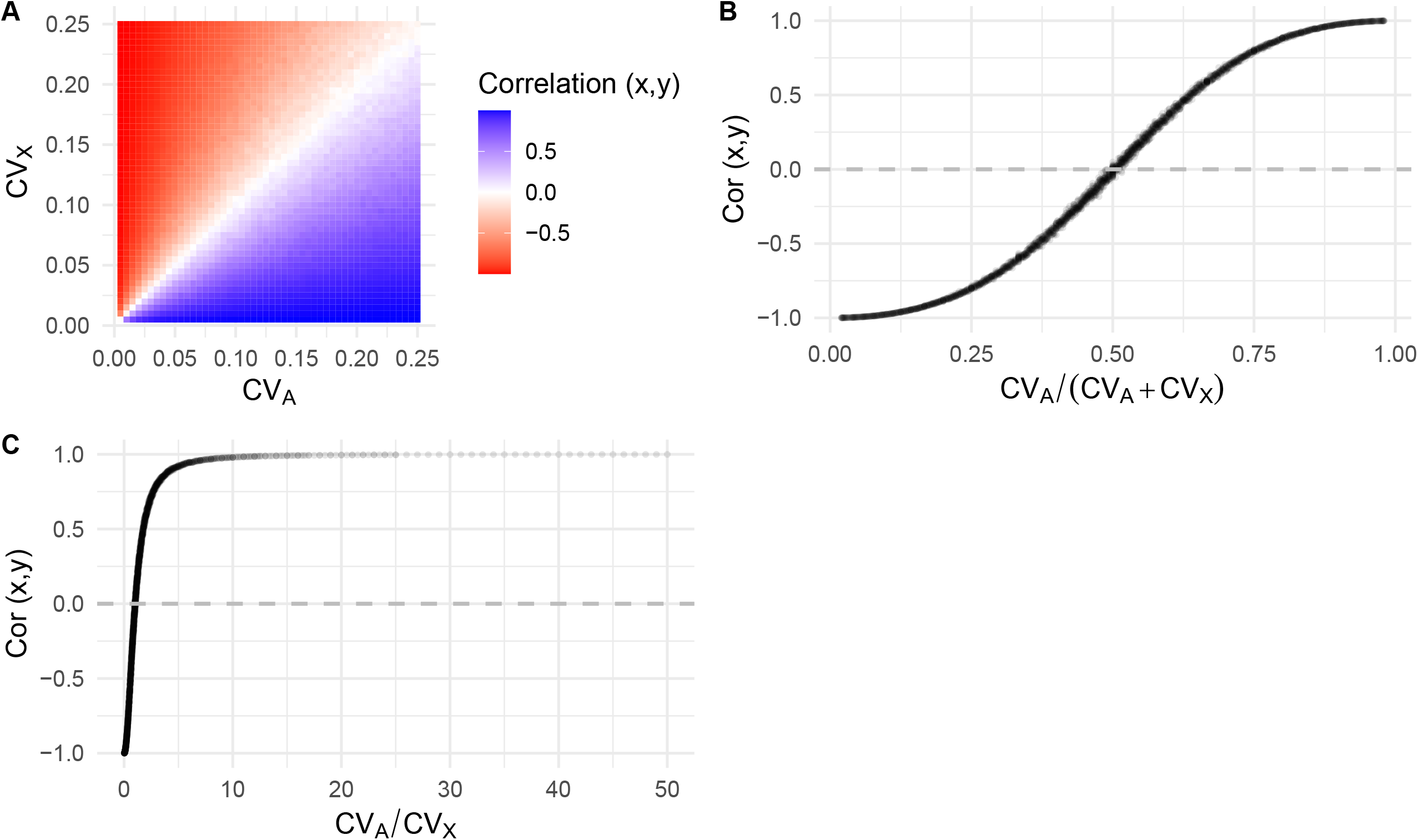
Model 1 results show a combined influence of CV_*A*_ and CV_*X*_, on the phenotypic correlation coefficient between traits *x* and *y* i.e. Cor(*x,y*). When CV_*A*_ = CV_*X*_, Cor(*x,y*) = 0; CV_*A*_ > CV_*X*_, Cor(*x,y*) > 0; CV_*A*_ < CV_*X*_, Cor(*x,y*) < 0. **(A)** Cor(*x,y*) represented as a heatmap for each combination of CV_*A*_ and CV_*X*_ values. **(B)** Cor(*x,y*) represented as a function of CV_*A*_/(CV_*A*_ + CV_*X*_), and of **(C)** CV_*A*_/ CV_*X*_. In **(A)**, each cell, and in **(B)** and **(C)**, each dot, represents 2500 simulated individuals.

### Aim 2

In model 2A and 2B, in addition to investigating the influence of CV_*A*_ and CV_*X*_, we tested whether μ_*A*_ or μ_*X*_ modulated Cor(*x,y*). Surprisingly, μ_*A*_ did not influence the phenotypic correlation between traits *x* and *y* (Figure 2A, 2B). The zero-crossing point, ‘*θ*_0_’, which is the value of ‘CV_*A*_/(CV_*A*_ + CV_*X*_)’ where Cor(*x,y*) changed sign, did not depend on μ_*A*_. In contrast, in model 2B, μ_*X*_ substantially modulated the influence of CV_*A*_ and CV_*X*_ on the phenotypic correlation between traits *x* and *y* (Figure 3A, 3B). Specifically, when allocation was less biased towards a trait (i.e. *X* < 0.5 < *Y*), ‘*θ*_0_’ was lower such that even when CV_*A*_ < CV_*X*_, positive phenotypic correlations occurred (Figure 3A, 3B, 3C). The opposite was true when *X* > 0.5 > *Y*. Whether *X* or *Y* were modelled as proportion allocation was inconsequential, because interchanging them had symmetrical meaning as long as the corresponding CV was used in the function (Figure 3D, also see model 4). Like results in model 1, in models 2A and 2B, relatively increasing CV_*A*_ led to a more positive phenotypic correlation, while relatively increasing CV_*X*_ led to a more negative correlation, between *x* and *y*.

**Figure 2:**
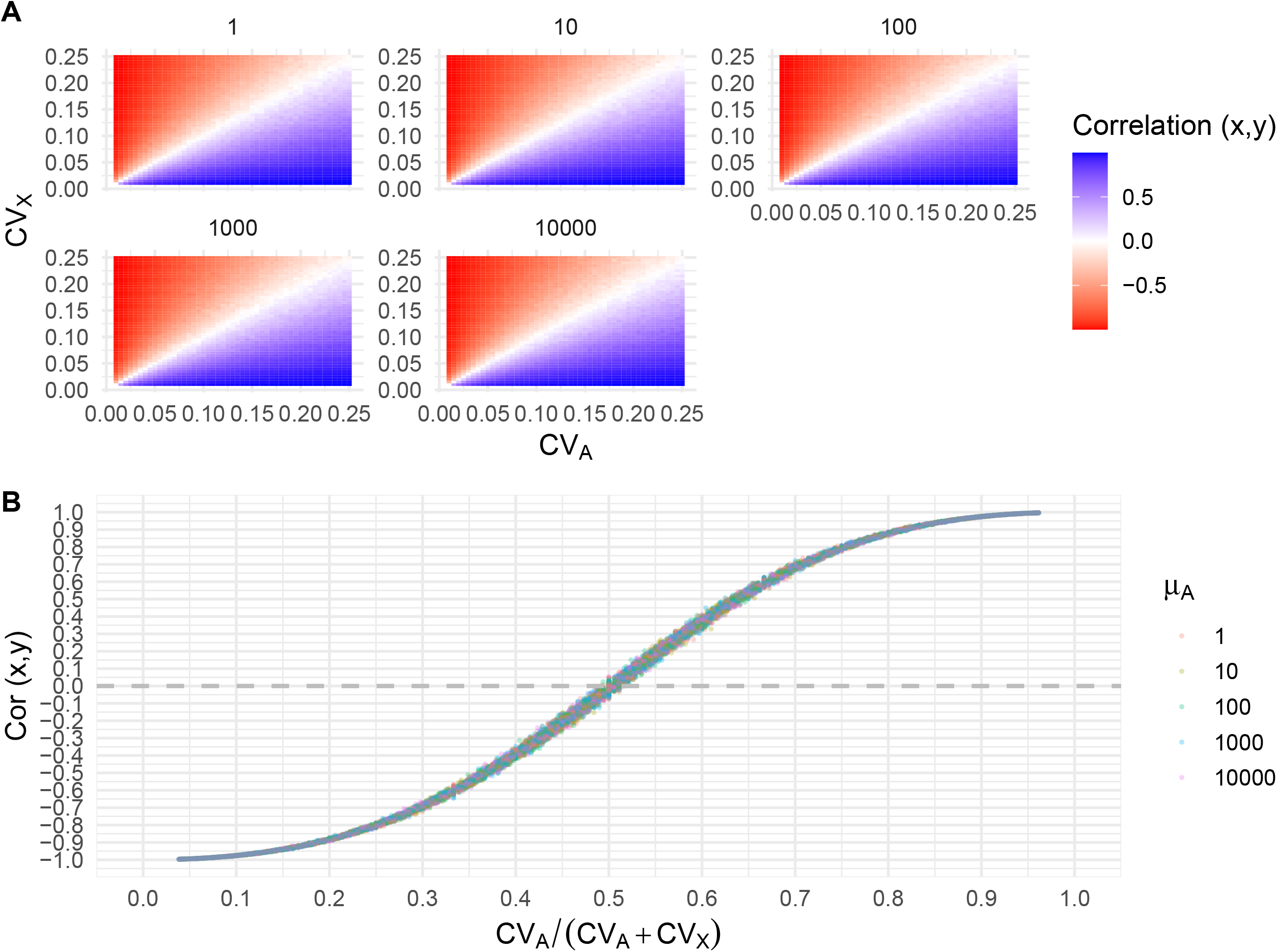
Model 2A results show a combined influence of CV_*A*_ and CV_*X*_ on Cor(*x,y*), but no influence of mean acquisition (μ_*A*_). **(A)** Cor(*x,y*) values do not differ between facets, each facet representing a different value of μ_*A*_. **(B)** Cor(*x,y*) is a function of CV_*A*_/(CV_*A*_ + CV_*X*_) but this dependency is not further modulated by μ_*A*_. Colours of dots represent a different μ_*A*_ values. In **(A)**, each cell, and in **(B)**, each dot, represents 2500 simulated individuals.

**Figure 3:**
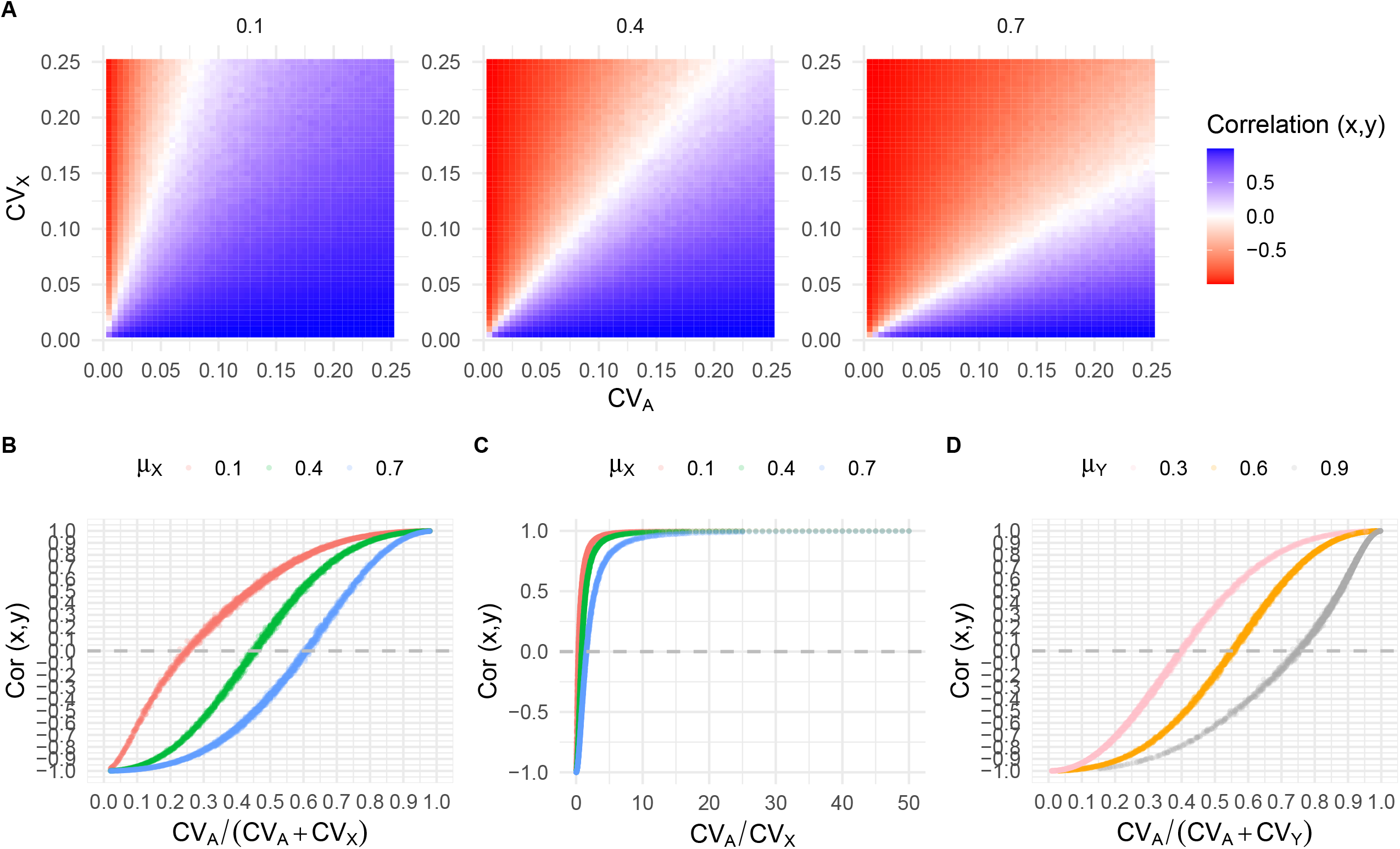
Model 2B results showing the combined influence of CV_*A*_ and CV_*X*_ on Cor(*x,y*), for different values of mean allocation strategy (μ_*X*_). **(A)** Cor(*x,y*) values in the heatmap differ between facets, each facet representing a different value of μ_*X*_, suggesting a modulating influence of mean allocation. **(B)** Cor(*x,y*) is a function of CV_*A*_/(CV_*A*_ + CV_*X*_) and this is further modulated by μ_*X*_. **(C)** Cor(*x,y*) is a function of CV_*A*_/CV_*X*_ and this is further modulated by μ_*X*_. **(D)** Cor(*x,y*) is a function of CV_*A*_/(CV_*A*_ + CV_*Y*_) and this is further modulated by μ_*Y*_. In **(A)**, each cell, and in **(B), (C)**, and **(D)**, each dot, represents 2500 simulated individuals.

### Aim 3

In model 3, we isolated the effects of CV_*A*_, CV_*X*_, μ_*A*_, an μ_*X*_ from effects of the covariance between acquisition and allocation, i.e. Cor(*A,X*) = 1 or Cor(*A,Y*) = -1, on the phenotypic trade-off between traits *x* and *y*. A positive correlation between acquisition (*A*) and allocation to trait *x* (i.e. *X*) intensified Cor(*x,y*), such that it went from negative to positive over smaller relative change in CV_*A*_ or CV_*X*_ (Figure 4, 5). This was because *x*_*i*_ = *X*_*i*_ · *A*_*i*_. Thus, when *X*_*i*_ had small values, *A*_*i*_ did too, thus *x*_*i*_ remains small; however, their large values exacerbated increases in *x*_*i*_ and amplified decreases in *y*_*i*_. In 3A and 3B, the shape of the function describing Cor(*x, y*) ∼ CV_*A*_/(CV_*A*_ + CV_*X*_) was substantially different compared to that in models 1, 2A, and 2B.

**Figure 4:**
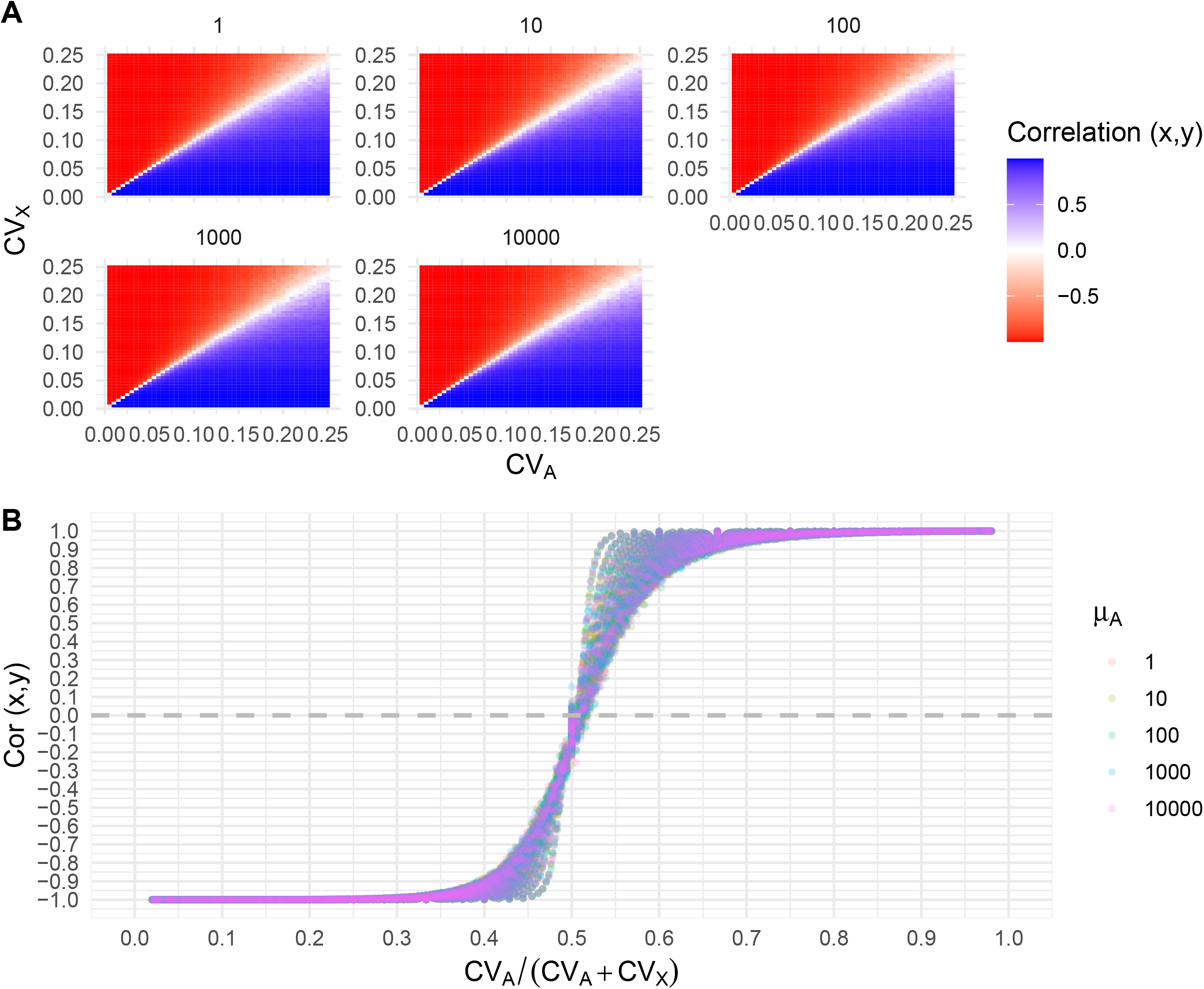
Model 3A results show a combined influence of CV_*A*_ and CV_*X*_, on Cor(*x,y*), but no influence of mean acquisition (μ_*A*_), when allocation and acquisition co-vary (Cor(*A,X*) = 1, Cor(*A,Y*) = -1). **(A)** Cor(*x,y*) values in the heatmap do not differ between facets, each facet representing a different value of μ_*A*_. However, compared to Figure 2A, the heatmap is darker, representing stronger strength of phenotypic correlations. **(B)** Cor(*x,y*) is a function of CV_*A*_/(CV_*A*_ + CV_*X*_) but this is not modulated by μ_*A*_. The shape of this function is different to the function in Figure 2B. Colours of dots represent different μ_*A*_ values. In **(A)**, each cell, and in **(B)**, each dot, represents 2500 simulated individuals.

**Figure 5:**
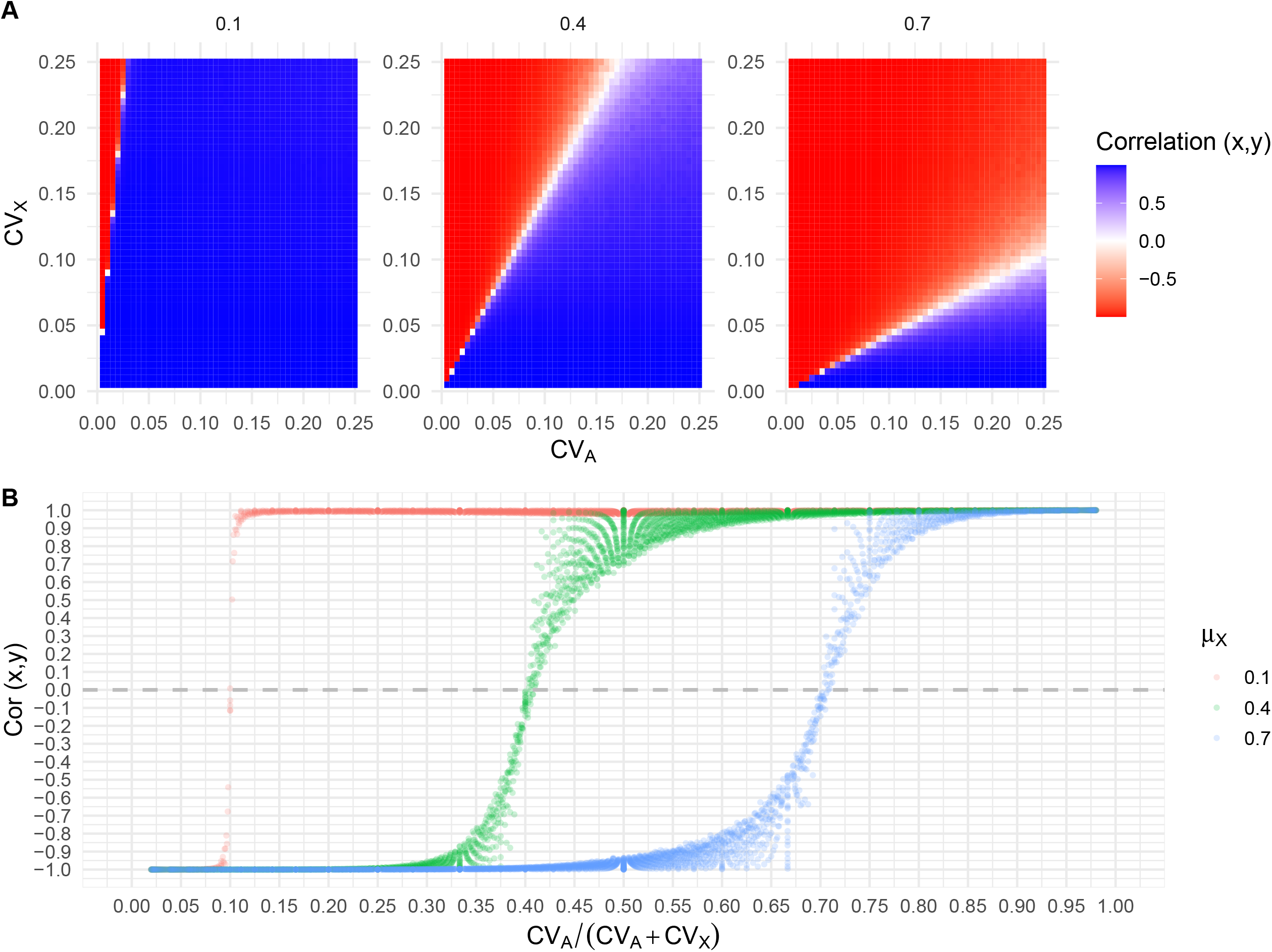
Model 3B results showing the combined influence of CV_*A*_ and CV_*X*_ on Cor(*x,y*), for different values of mean allocation strategy (μ_*X*_) when allocation and acquisition co-vary (Cor(*A,X*) = 1, Cor(*A,Y*) = -1. **(A)** Cor(*x,y*) values in the heatmap differ between facets, each facet representing a different value of μ_*X*_, suggesting a modulating influence of μ_*X*_. However, compared to Figure 3A, the heatmap is darker due to higher magnitude of phenotypic correlations. **(B)** Phenotypic correlations between traits *x* and *y* are a function of CV_*A*_/(CV_*A*_ + CV_*X*_) which is further modulated by μ_*X*_. Compared to Figure 3B where *A* and *X* do not co-vary, here, positive covariance between *A* and *X* creates a steeper function, with *θ*_0_ being the same as μ_*X*_.

Like model 2A, in 3A, μ_*A*_ did not modulate the influence of CV_*A*_, CV_*X*_ on Cor(*x,y*). However, like model 2B, in 3B, μ_*X*_ modulated this relationship (Figure 4, 5). However, in 3B, *θ*_0_ was substantially more affected by μ_*X*_ compared to in 2B (i.e. Figure 5B and 5C versus Figure 3A). Specifically, in 3B, *θ*_0_ had the same value as μ_*X*_ (Figure 5B, 5D). Cor(*A,X*) = -1, led to identical results as Cor(*A,Y*) = 1, except here, the modulating influence was of μ_*Y*_ and CV_*Y*_ (Supplementary section S1; Figure S23).

### Statistically quantifying effects

In model 4 where all parameters were varied simultaneously, the linear model that best described the data contained the terms (Table 2): CV_*A*_/(CV_*A*_ + CV_*X*_): μ_*X*_: Cor(*A, X*) + CV_*A*_/(CV_*A*_ + CV_*Y*_). In this model, all terms had a significant influence (P < 0.001) on Cor(*x,y*) (Figure S23) and collectively explained 82.6% of the variation in phenotypic correlations. μ_*A*_, whenever present in any of the other linear models, had no significant influence (P > 0.9) on phenotypic correlations (Table S3). Importantly, the ratio ‘CV_*A*_/(CV_*A*_ + CV_*X*_)’ yielded a better fitting model than the interaction ‘CV_*A*_: CV_*X*_’, or the ratio ‘CV_*A*_/CV_*X*_’ (Table 2). These statistically reinforce the results of models 1-3.

### Validation with empirical data

We used empirical data on the parameters CV_*A*_, CV_*X*_, μ_*A*_, μ_*X*_, and Cor(*A,X*) from King et al (2011) to simulate a population of individuals with two traits and calculate their among-individual phenotypic correlation. The simulated phenotypic correlations matched those reported by King et al (2011b) (R^2^ = 0.9896; Supplementary section 3, Figure S22).

Individually changing the empirically observed μ_*A*_ value had no influence on phenotypic correlations. However, individually changing the empirical values of any other parameter, i.e. of CV_*A*_, CV_*X*_, μ_*A*_, μ_*X*_, and Cor(*A,X*) altered the simulated phenotypic correlation.

## Discussion

### Summary

The Y-model is a heuristic for why among-individual phenotypic correlations between life-history traits are positive, despite expected trade-offs. It offers a fundamental biological and statistical understanding of the detectability of trade-offs. However, the Y-model does not make *a priori* quantitative predictions about the magnitude of trade-offs or clarify the mechanisms by which these are modulated. It is also unclear how key parameters of the model, namely, means of acquisition and allocation, their covariance, and their variance, interact to influence phenotypic correlations. In line with previous studies (Descamps et al, 2016; Johnson and Nasrullah, 2024; Roff and Fairbairn, 2007; Robinson and Beckerman, 2013; Van Noordwijk and De Jong, 1986), we find that the relative coefficients of variation of acquisition and allocation influence the detection of phenotypic trade-offs (aim 1). However, we show that this is modulated by the mean allocation strategy of the population, but surprisingly, not by the mean resources acquired by a population (aim 2). Finally, positive or negative covariances between acquisition and allocation exacerbate the strength of phenotypic correlations (aim 3), and this effect is further influenced by mean allocation but not by mean acquisition. The mechanisms of phenotypic correlation in the Y-model can thus be expressed as (Figure S23, Table 2, S2, S3):

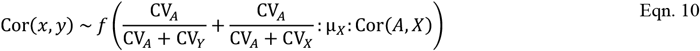

### Dietary restriction, resource richness, and mean acquisition

Populations experiencing dietary abundance or in resource rich environments often do not show among-individual phenotypic trade-offs, while populations on restricted diets do (e.g. Adler et al, 2013; Attisano et al, 2012; Collins et al, 2023; Messina and Fry, 2003; Richerdson and Smiseth, 2020; Robert et al; 2015; Smiseth et al, 2014; Tatar and Carey, 1995; Treidel et al, 2021). Among other explanations (Irish et al, 2025), the Y-model is used to rationalise this result. But this is a misinterpretation of mechanisms in the Y-model that modulate trade-offs. The model does not predict that increasing resources available to a population on average (μ_*A*_) masks trade-offs while decreasing μ_*A*_ reveals trade-offs. Instead, it states that increase in the relative variation in acquisition compared to allocation strategy masks trade-offs (van Noordwijk and De Jong, 1986; Mogensen and Post, 2012; Robinson and Beckerman, 2013). This is supported by our simulations. Therefore, altering the average resource richness of an environment (e.g. dietary restriction) would have no direct effect on phenotypic correlations when CV_*A*_, CV_*X*_, Cor(*A,X*), and μ_*X*_ are held constant. Here, scaling each individual’s resource (*A*_*i*_) by a factor ‘*c*’ when the population-level diet (μ_*A*_) is improved by that factor (*c*μ_*A*_), leads to the phenotypic values (*x* and *y*) and their standard deviations being similarly scaled, such that Cor(*cx,cy*) *=* Cor(*x,y*).

We hypothesize that the oft-observed effect of resource restriction on trade-offs (e.g. Richardson and Smiseth, 2020) are not directly driven by changes in population-level mean acquisition (μ_*A*_) (‘mean invariance hypothesis’, Figure 6). Instead, we propose three alternative testable predictions for mechanisms modulating phenotypic trade-offs when the resource available to a population of individuals is restricted or increased. First, between-individual differences in the ability to process food or convert food to energy (e.g. Olijnyk and Nelson, 2013; reviewed in Boggs, 1992; Roff and Fairbairn, 2007) could lead to among-individual differences in the “ceilings” of resources that can be potentially acquired (Figure 6A). This would cause dietary restriction to reduce CV_*A*_ thereby revealing phenotypic trade-offs, while dietary abundance increasing CV_*A*_ and masking trade-offs (Figure 6A). Second, increase or decrease in mean resource abundance might change how individuals are allocating resources (De Lisle and Rowe, 2023). For example, under restricted diets, individuals might allocate proportionately less resources to reproduction and proportionately more to survival (Aronoff and Trumble, 2025; Moat et al, 2016; Shanley and Kirkwood, 2000); or employ the opposite allocation strategy, as seen under terminal investment (Krams et al, 2015). This could be a consequence of either allocation to one trait being prioritised over another, or traits having different ceilings for energy that can be invested in them due to physiological constraints (Figure 6B). Thus, dietary restriction could alter the covariance between acquisition and allocation strategy, Cor(*A,X*), thereby altering phenotypic trade-offs (e.g. King et al, 2011a). Third, dietary restriction might impact the mean allocation strategy (μ_*X*_) or the variation in allocation (CV_*X*_) of a population (e.g. Saeki and Crowley, 2013; Wang et al, 2025). This could happen via genotype by diet interactions (Figure 6C; Geberhardt and Stearns, 1988, 1993a, 1993b; Flatt, 2014; Kent et al, 2009) where different genotypes vary in their reaction norms (e.g. De Lisle and Rowe, 2023; Zajitschek and Connallon, 2017). The mean invariance hypothesis provides precise, testable predictions (Figure 6) for which distinct mechanisms reveal phenotypic trade-offs in resource limited environments, thus clarifying previous misunderstandings of the Y-model. Future studies could extend this hypothesis’ scope, by exploring if dissimilar phenotypic trade-offs between two populations that differ in their environmental conditions (e.g. temperature, predation), is caused due to differences in μ_*X*_, CV_*X*_, CV_*A*_, or Cor(*A,X*) between these populations.

**Figure 6:**
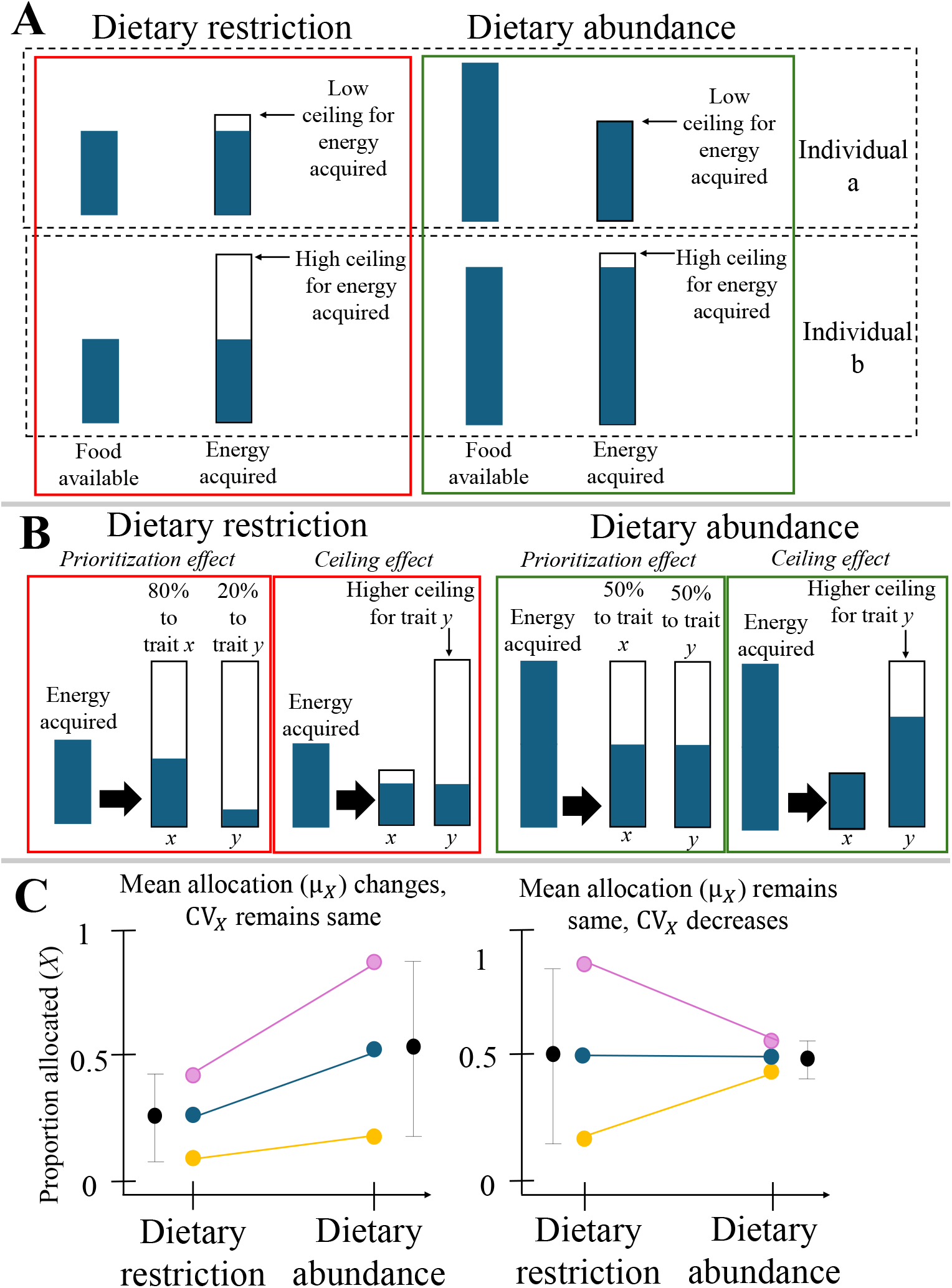
Conceptual diagram of the ‘mean invariance hypothesis’ and its associated predictions for the mechanisms modulating phenotypic trade-offs under dietary abundance. **(A)** Dietary restriction changes phenotypic correlations between traits *x* and *y* by changing CV_*A*_ due to among-individual differences in ceilings for acquisition. Filled (blue) quantity shows the absolute amount of energy acquired from food, empty (white) quantity shows the amount that can still be acquired due to the ceiling not being reached. **(B)** Dietary restriction changes phenotypic correlation between traits *x* and *y* due to covariance between *A* (acquisition) and *X* (proportion allocation). Under restricted diets, one trait might be prioritised and proportionately more resources are allocated towards it (i.e. higher *X*) compared to the other trait, while under abundant diet proportionately less resources are allocated towards it (i.e. higher *Y*). Alternatively, one trait (trait *x*) might have a lower ceiling than the other, therefore under restricted diets, similar absolute energy is invested in traits *x* and *y*, but under abundant diets, trait *y* get more energy because its ceiling has not been reached yet. **(C)** Dietary restriction changes phenotypic correlation between traits due to among-individual differences in reaction norms. Diet either alters mean allocation strategy of a population (μ_*X*_) (left panel) or the coefficient of variation in allocation (CV_*X*_) (right panel). Each colour shows one individual. Black dot and error bars show hypothetical mean and variance (σ^2^) of the sampled population.

### Evolution of trade-offs

Phenotypic trade-offs are a key component of evolutionary theory as they apply constraints on phenotypic-space (Cohen et al, 2019) and determine whether selection can directionally improve a trait (Kingsolver and Diamond, 2011; Roff and Fairbairn, 2007). In addition to reflecting resource limitation (as formalised in the Y-model), phenotypic trade-offs can also be a consequence of physiological constraints or antagonistic pleiotropy (Chang et al, 2024; Garland et al, 2022; Harshman and Zera, 2007; Spitze et al, 1991). These are not mutually exclusive, because variation in allocation strategy could be underpinned by heritable genetic variation (Robinson and Beckerman, 2013). If so, the mean allocation strategy of a population (μ_*X*_) will represent its current evolutionary state, while the covariance between acquisition and allocation could represent underlying genetic pleiotropy or linkage disequilibrium (Sgrò & Hoffmann, 2004). Like others, we emphasize that the absence of phenotypic trade-offs does not imply absence of genetic trade-offs (Haave-Audet, 2022; De Laguerie; Robinson and Beckerman, 2013; Roff and Fairbairn, 2007), and caution empiricists to not conflate the two.

The distinction between genetic versus phenotypic trade-offs becomes crucial when studying senescence (Boggs, 2009; Cohen et al, 2020; Lemaitre et al, 2015; Maklakov et al, 2017; Maklakov and Chapman, 2019; Takeshita, 2024). Due to a weaker selection in late-life, genes with antagonistic effects that improve early-life reproduction at the cost of lowering late-life reproduction or survival, are positively selected for (Maklakov and Chapman, 2019). Although contrary to predictions, studies often find that individuals with higher early-life fecundity also have higher late-life fecundity, or higher survival (e.g. Bouwhuis et al, 2009; Jehan et al, 2022; Sanghvi et al, 2025a; Sanghvi et al, 2024; Van De Pol and Verhulst, 2006; Winder et al, 2025). This relation, however, can occur despite within-individual genetic pleiotropy that determines partitioning of resources to different traits (Haave-Audet et al, 2022). Furthermore, ontogenic and age-specific variation in allocation strategy and resource acquisition (Barreaux et al, 2022; De Jong, 1993; Garland et al, 2022; Harshman and Zera, 2007; Heino and Kaitala, 1999; Worley et al, 2003; Zera and Harshman, 2001), as well as allocation-acquisition autocorrelation between different ages, could lead to phenotypic trade-offs being dependent on organismal age (Davison et al, 2014; Van Den Heuvel et al, 2017). Theoreticians could incorporate temporality in our quantitative models to refine theories for how ageing impacts phenotypic trade-offs.

### Pace-of-life, polymorphism, plasticity

Differences in allocation of resources are likely to occur along the slow-to fast-pace-of-life syndrome (“POLS”) axis (Healy et al, 2019; Laskowski et al, 2021; Montiglio et al, 2018; Salzman et al, 2018; Stott et al, 2024). After accounting for effects of phylogeny, our models could be used to generate specific predictions for phenotypic trade-offs along the POLS axis (also see Descamps et al, 2016). ‘Fast-lived’ species that bias allocation towards early-life reproduction (μ_*X*_ > 0.5) might be more likely to show phenotypic trade-offs between early-life fecundity and survival, than ‘slow-lived’ species. This is because the zero-crossing point (*θ*_0_) in fast-living organisms occurs at higher values of ‘CV_*A*_/(CV_*A*_ + CV_*X*_)’ compared to organisms with a slow pace-of-life (‘allocation bias hypothesis’; Figure 7). These patterns would be further exacerbated when there is positive covariance between acquisition and allocation to early-life fecundity, but will be buffered when there is negative correlation between these (‘covariance amplifier hypothesis’; Figure 7, S23).

**Figure 7:**
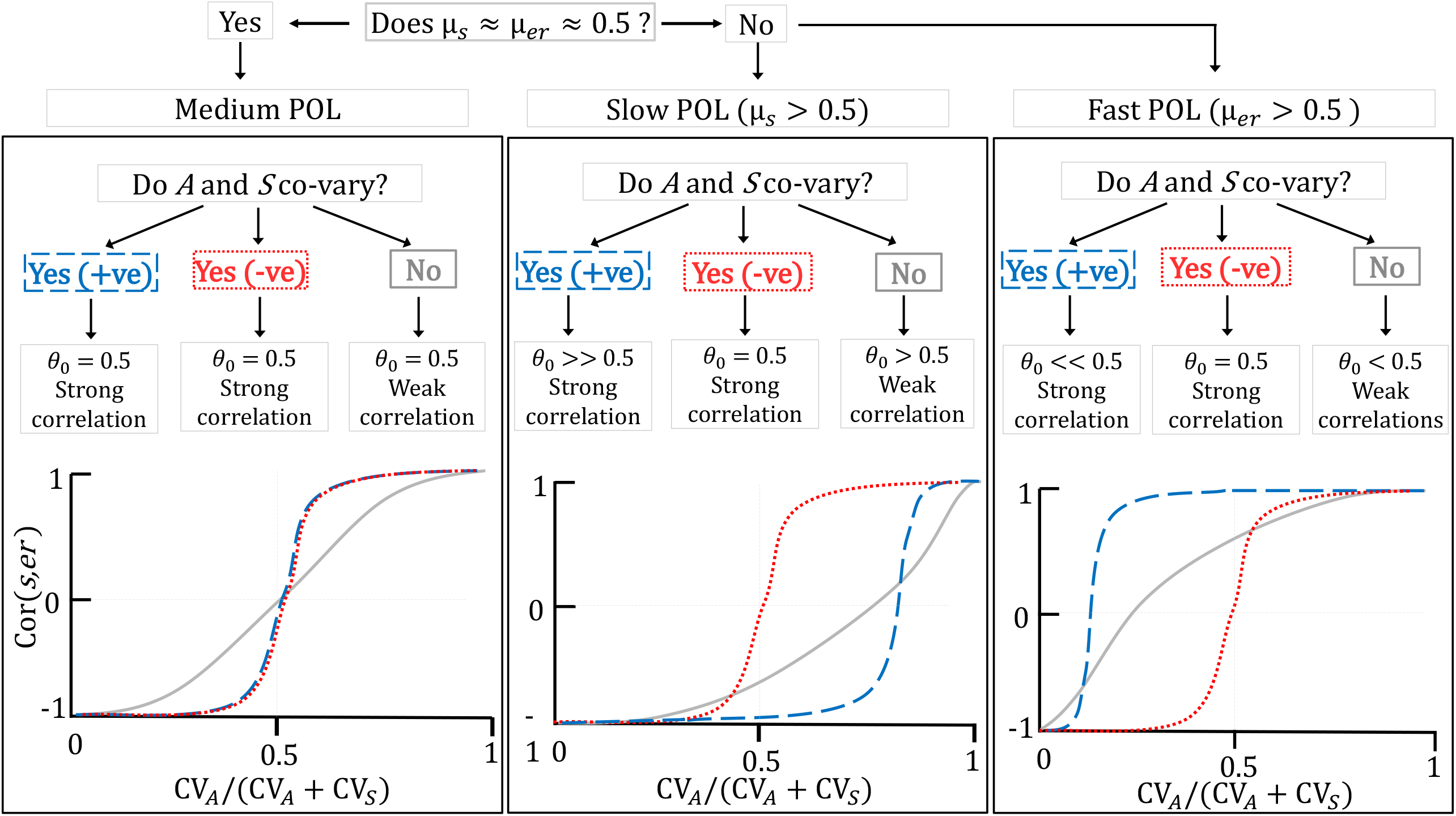
Conceptual diagram summarizing the ‘allocation bias’ and ‘covariance amplifier’ hypotheses. ‘Allocation bias hypothesis’ compares slow vs. fast POLS, while ‘covariance amplifier hypothesis’ compares the effects of negative, zero, or positive covariance between allocation on acquisition. μ_*s*_ and μ_*er*_ show the average resource allocation strategy of a population towards survival and early-life reproduction, respectively. These two traits are chosen as hypothetical biological examples because allocation strategy towards these corresponds well to a population’s POLS. CV_*S*_ is the CV of proportion resources allocated to survival; CV_*A*_ is the CV of acquisition. Grey, red, and blue represent zero, negative, and positive covariance between *A* and *S*, respectively, that is the within-individual covariance between acquisition and proportion allocation towards survival. ‘*θ*_0_’ is the zero-crossing point, i.e. the value of CV_*A*_/(CV_*A*_ + CV_*X*_) where the phenotypic correlation, Cor(*s,r*), changes sign.

Our Y-model generates some predictions for how trade-offs are impacted by alternative morphs (e.g. Stec et al, 2025). In species where alternative morphs allocate proportionately different fractions of resources towards different traits (Miles et al, 2007; Oliveira et al, 2008), variation in allocation would be relatively higher compared to species without such polymorphism. Thus, phenotypic trade-off in polymorphic species might be easier to detect. However, these trade-offs could be modulated if polymorphism (i.e. resource allocation strategy) is condition-dependent (Buzzato et al, 2011; De Lisle and Rowe, 2023; Engqvist and Taborsky, 2015; McNamara and Houston, 1996; Ng’oma et al, 2017; Roff, 2011; Smallegange et al, 2012). For example, if a male on abundant resources develops into a morph that invests heavily in intra-sexual competition (i.e. Cor(*A,X*) → Cor(*diet, competition*) > 0, and μ_*competition*_ > 0.5) and less in sperm numbers (Knell and Parrett, 2024; Sanghvi et al, 2025b), within this morph, phenotypic trade-offs between intra-sexual competition-related traits and sperm numbers would be amplified in magnitude (‘covariance amplifier hypothesis’). Here, phenotypic trade-offs would also be easier to detect due to *θ*_0_ occurring at larger values of CV_*A*_/(CV_*A*_ + CV_*X*_) (Figure 7).

Covariance between resource acquisition and allocation strategy has important consequences for the evolution of trade-offs over time (King and Roff, 2010). In stable environments, plasticity in life-history strategies is hypothesized to reduce, while in predictably fluctuating environments, selection favours plastic strategies and steeper reaction norms (DeWitt et al, 1998; King and Hadfield, 2019; Murren et al, 2015; Nussey et al, 2007; Vinton et al, 2022). Therefore, in stable environments, the plastic dependence of life-history strategy on resource acquisition, i.e. Cor(*A,X*), would be closer to zero, compared to fluctuating environments. This provides a possible mechanism for why over time, environmental stability buffers phenotypic trade-offs while environmental fluctuation exacerbates them (King and Roff, 2011). Furthermore, if selection acts on reaction norms (Brommer et al, 2007; Kingsolver et al, 2006; Murren et al, 2014; Van Tienderen and Koelewijn, 1994), over time this could alter μ_*X*_ and reduce CV_*X*_ (Figure 6C), therefore changing phenotypic trade-offs in populations (e.g. Rogissart et al, 2025; Roff et al, 2002).

The often-observed lack of phenotypic trade-offs (Have-Audet et al, 2022; Winder et al, 2025) can be explained by other mechanisms in the Y-model. If allocation strategy among individuals in a population is homogeneous because of canalisation (Gibson and Wagner, 2000; Kawecki, 2000; Mieklejohn and Hartl, 2002) or past episodes of intense selection, detection of phenotypic trade-offs will be less likely. Generally, greater statistical potential for variation in acquisition, than allocation which is bounded between zero and one (Morris and Doak, 2004), could also lead to a low tendency for phenotypic trade-offs being detected across taxa. Finally, in species where acquisition is highly variable, for example, where foraging patches differ [randomly] in resource abundance or individuals differ substantially in foraging efficiency (Boggs, 1992; Pontzer and McGrosky, 2022), phenotypic trade-offs might be masked (Robinson and Beckerman, 2013; Reznick et al, 2000).

### Testing hypotheses

Our extension of the Y-model generates some novel predictions for the presence or absence of phenotypic trade-offs, which could be empirically tested by future studies. Empirical tests of the Y-model in the past have been limited, due to the mechanisms creating phenotypic correlations being a black box (reviewed in Hashemi et al, 2024; Laskowski et al, 2021; Roff and Fairbairn, 2007). Additionally, the simplicity of Y-model assumptions (described in Descamps et al, 2016; Johnson and Nasrullah, 2024; Roff and Fairbairn, 2007; Robinson and Beckerman, 2013) have hindered its empirical applicability. For example, allocation of resources to only two traits or from a common resource pool does not represent biological reality (De Jong, 1993; Garland et al, 2022; Pease and Bull, 1988; Worley et al, 2003).

Similarly, assuming that allocation and acquisition are sampled from normal distributions (Johnson and Nasrullah, 2024), there is perfect conversion of resources to energy (Boggs, 1992; Roff and Fairbairn, 2007), or traits use the same currencies (Cohen et al, 2017), can be unrealistic. Furthermore, Y-models often assume a lack of covariance between allocation and acquisition (e.g. Van Noordwijk and De Jong, 1986; Roff and Fairbairn, 2007); however, such covariances are common (King et al, 2011a, 2011b) and can alter predictions (Descamps et al, 2016; Robinson and Beckerman, 2013; Zajitschek and Connallon, 2017; our results). We second these criticisms and strongly caution empiricists to beware of their assumptions when using this framework. We also take specific steps to address some of these past limitations. For example, we conduct sensitivity analyses with complex distributions to ensure robustness of models; simulate covariance between allocation and acquisition; validate our models with empirical data; and importantly, systematically partition various interacting mechanisms to reveal key pathways in the “black box”. Despite its simplicity, we emphasize this framework’s past success as a heuristic, and through our study, as a tool with quantitative predictions, to understand trade-offs. While Pelabon et al (2020) have been critical of using CV to quantify trait variation, we highlight its usefulness when used in biologically meaningful contexts and in relation to other parameters. Future work could derive equation 10 analytically (see Supplementary section S4) and add complexity to our quantitative framework by incorporating more than two traits, different currencies and pools for allocation (Cohen et al, 2017), and imperfect resource conversion. While the Y-model is typically used for understanding among-individual phenotypic correlations, when repeated measures of two traits are available for an individual (e.g. at different ages), the model can also be used to understand phenotypic correlations within-individuals.

An overlooked way to test Y-model predictions involves using simple systems where the entire acquired resource pool is allocated towards only two components, and acquisition and allocation are measured easily, and importantly, in the same units. We highlight three such systems, and encourage researchers to look for more. First, studies that measure the numbers of sperm inseminated by males into females, the numbers of sperm ejected, and the numbers of sperm retained by females, could be used (Sanghvi et al, 2025c). Here, the total numbers of sperm inseminated can be modelled as resource acquisition, the total numbers ejected and retained as the phenotypic values of *x* and *y*, and the proportion ejected versus retained as the fractional allocation strategy, *X* and *Y*, respectively. By experimentally manipulating variation in the numbers of sperm inseminated (i.e. CV_*A*_) using lines selected for increased or decreased sperm ejection (i.e. manipulating μ_*X*_), or using populations with homogenous ejection strategies (i.e. manipulating CV_*X*_), the influence of different Y-model parameters can be tested in a biological setting. Second, studies where early- and late-life reproduction are assayed could be used. Here, total reproductive success (early + late) is resource acquisition, the fraction of offspring produced in early-versus late-life is the resource allocation strategy (*X* and *Y*), and the correlation between numbers of offspring produced early versus late is Cor(*x,y*). Third, studies on allometry, where sizes of two different body parts are measured (e.g. King et al, 2011a, 2011b; Meczek and Nijhout, 2004; Saglam et al, 2008) could be used. Here, total mass of studied organs represents acquisition, relative mass of each part represents allocation strategy, and the allometric slope of the mass of the two parts represents the phenotypic correlation. These systems might not be typically viewed as interesting representations of the acquisition-allocation Y-model. However, they are an ideal way to experimentally manipulate and test each parameter’s influence. Such systems also underscore the Y-model’s broad applicability beyond life-history theory, to explain covariances in any system structurally alike equations 1-3, such as the association between owning large houses and buying expensive cars (Reznick et al. 2000).

## Conclusions

The Y-model is typically used as a heuristic for explaining the presence or absence of phenotypic trade-offs. We extend it into a predictive, quantitative framework that partitions different components of allocation and acquisition, and pinpoints the mechanisms directly driving phenotypic correlations. We show that contrary to common misinterpretations of the model, phenotypic correlations are not directly dependent on mean resources acquired by populations, but rather, on the interaction between the variation in allocation and acquisition, mean allocation strategy, and covariance between allocation and acquisition. Importantly, our models generate three distinct hypotheses (Figure 6, 7). These outline specific predictions for phenotypic trade-offs in the context of pace-of-life syndromes, dietary restriction, and polymorphism. We demonstrate the robustness of our models, validate our predictions with empirical data, and highlight examples of simpler systems that can be used to test Y-model predictions. We caution that researchers do not conflate genetic and phenotypic trade-offs, and be aware of assumptions of the Y-model. Despite these assumptions, our systematic partitioning of various parameters (Figure 7; Figure S23) provides a powerful, quantitative tool to understand life-history evolution. Importantly, this reveals the minimum information required to accurately predict phenotypic trade-offs in systems where resources are acquired and allocated. Our study can guide empiricists in life-history research by detailing what they should be trying to measure and why; how they could build and test hypotheses; and how to interpret the presence or absence of phenotypic trade-offs.

## Acknowledgements

We are deeply grateful to Derek Roff for extensively debating these ideas with us, providing comments, analytical solutions and constructive criticism. We also thank Stephen Stearns and Rob Salguero-Gómez for commenting on the manuscript; David Reznick, Alex Kacelnik, Jeff Lemaitre, and Jessica Metcalfe for their valuable suggestions; and fly lab members for their feedback.

## Use of AI statement

DeepSeek R-1 and Gemini 2.5 were used to streamline the simulation code.

## Supporting information

### Supplementary figures

**Figure S1:**
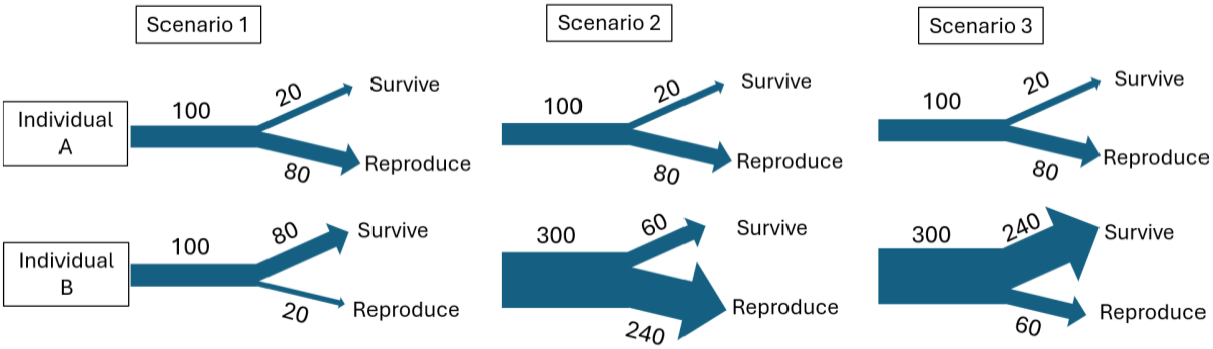
The Y-model represented as resources being acquired and subsequently allocated to two traits (e.g. survival and reproduction). Three hypothetical scenarios shown. In scenario 1, due to among-individual variation in allocation strategy but no variation in acquisition between individuals, the resultant phenotypic covariance between survival and reproduction will likely be negative. In scenario 2, due to variation in resource acquisition, but no variation in the proportion of resources allocated to survival and reproduction (i.e. allocation strategy), the resultant phenotypic covariance between survival and reproduction will likely be positive. In scenario 3, there is variation in both, resource allocation and acquisition.

**Figure S2:**
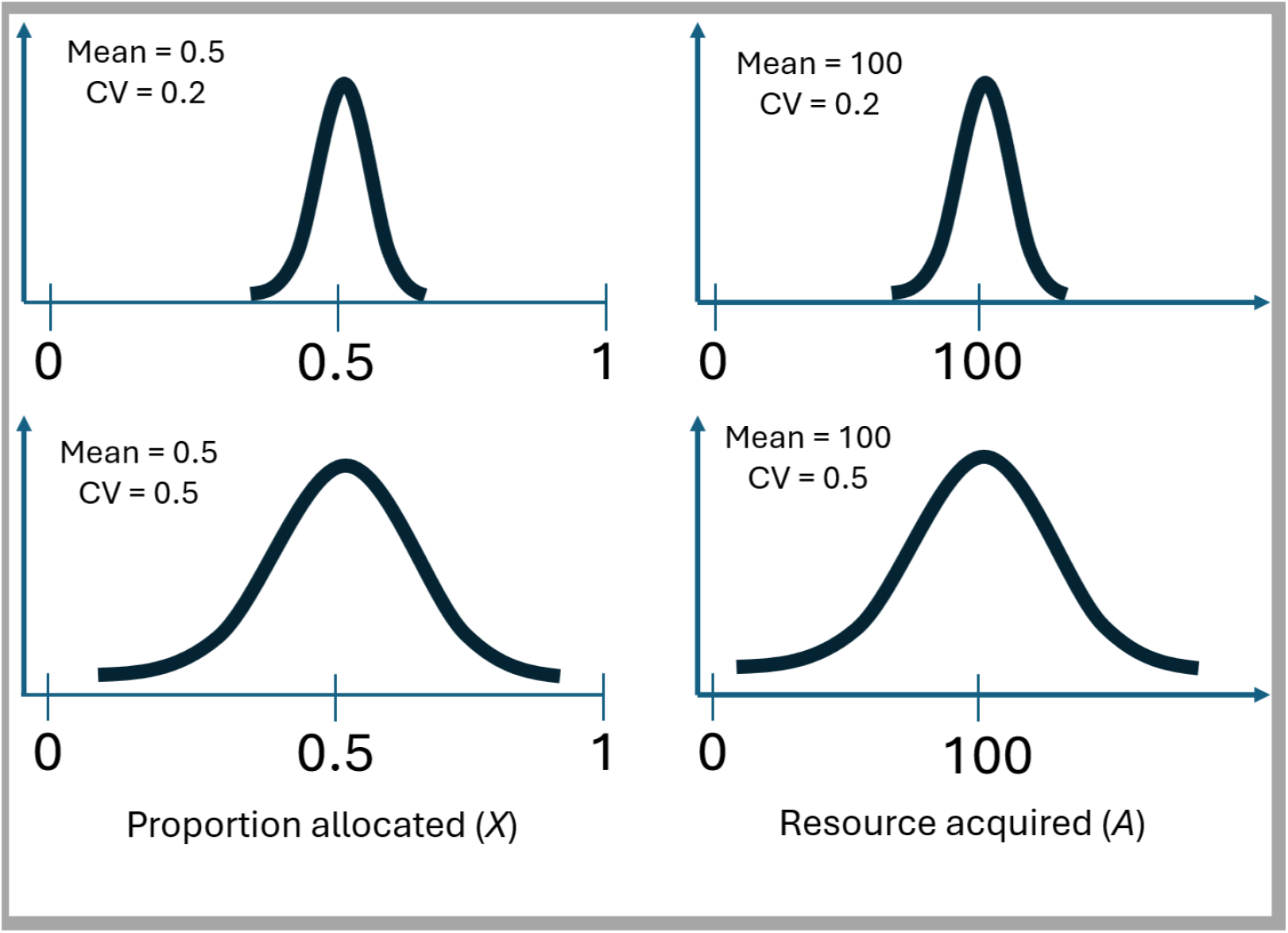
An example of the comparisons we simulated in aim 1, where we modelled different combinations of CV_*A*_ and CV_*X*_, while holding the means constant, to understand how CV impact the phenotypic correlation between traits *x* and *y*. Left panels represent an example of different CV_*X*_ values with the same μ_*X*_, right panels show an example of different CV_*A*_ with the same μ_*A*_.

**Figure S3:**
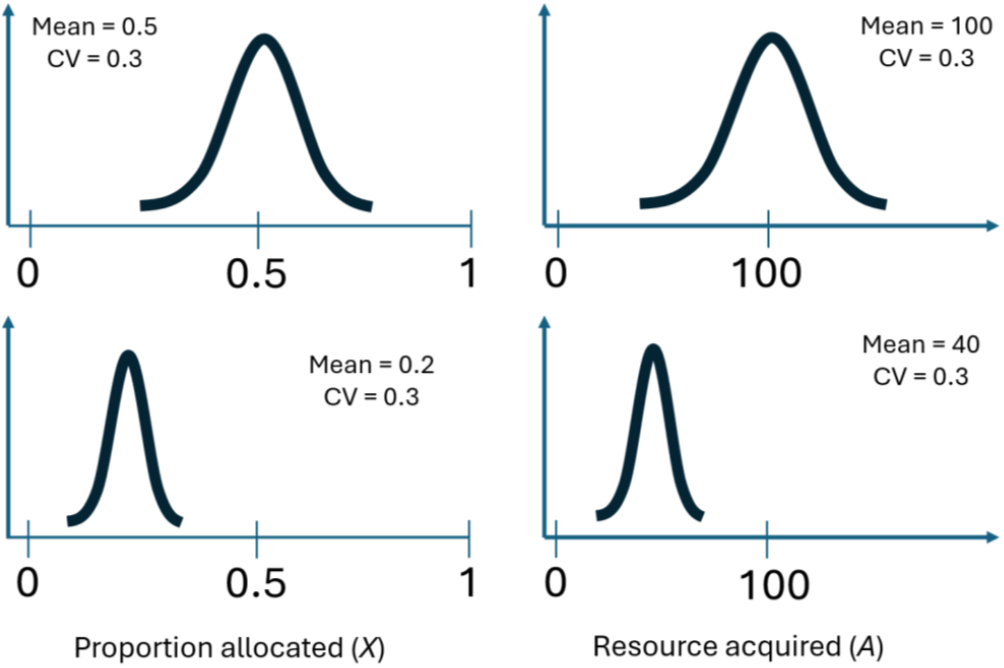
An example of the comparisons we simulated in model 2A and 2B, where not only different combinations of CV_*A*_ and CV_*X*_ were iterated (Figure S2), but also different values of μ_*X*_ or μ_*A*_ were, to understand how the means impact the phenotypic correlation between traits *x* and *y*. Left panels represent an example of same CV_*X*_ values with different μ_*X*_, right panels show an example of same CV_*A*_ with different μ_*A*_.

**Figure S4:**
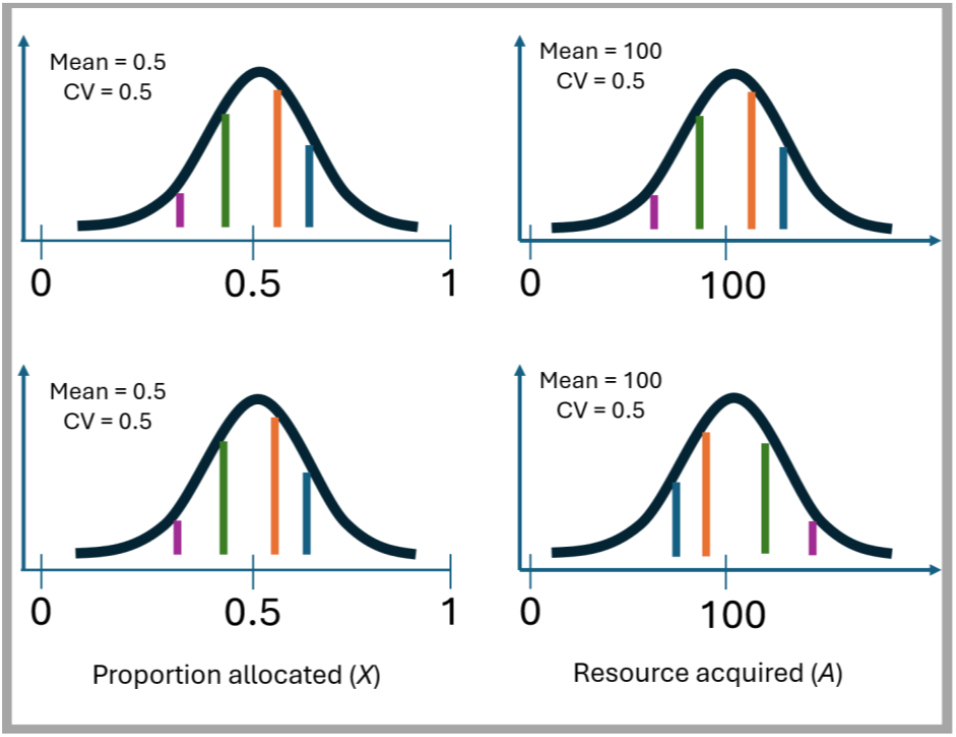
Hypothetical example of the comparisons in model 3, where we not only simulated different combinations of CV_*A*_ and CV_*X*_ (Figure S2) and iterated different values of μ_*X*_ or μ_*A*_ (Figure S3), but also co-varied *X* and *A* within-individuals, to understand how these impact the phenotypic correlation between traits *x* and *y*. Left and right panels represent distributions of *X* and *A* respectively, both with the same CV. Top panel shows positive covariance between *A*_*i*_ and *X*_*i*_, bottom panel shows negative covariance between *A*_*i*_ and *X*_*i*_. Each coloured line represents a set of different individuals and their relative position in the distribution of *A* and *X*.

**Figure S5:**
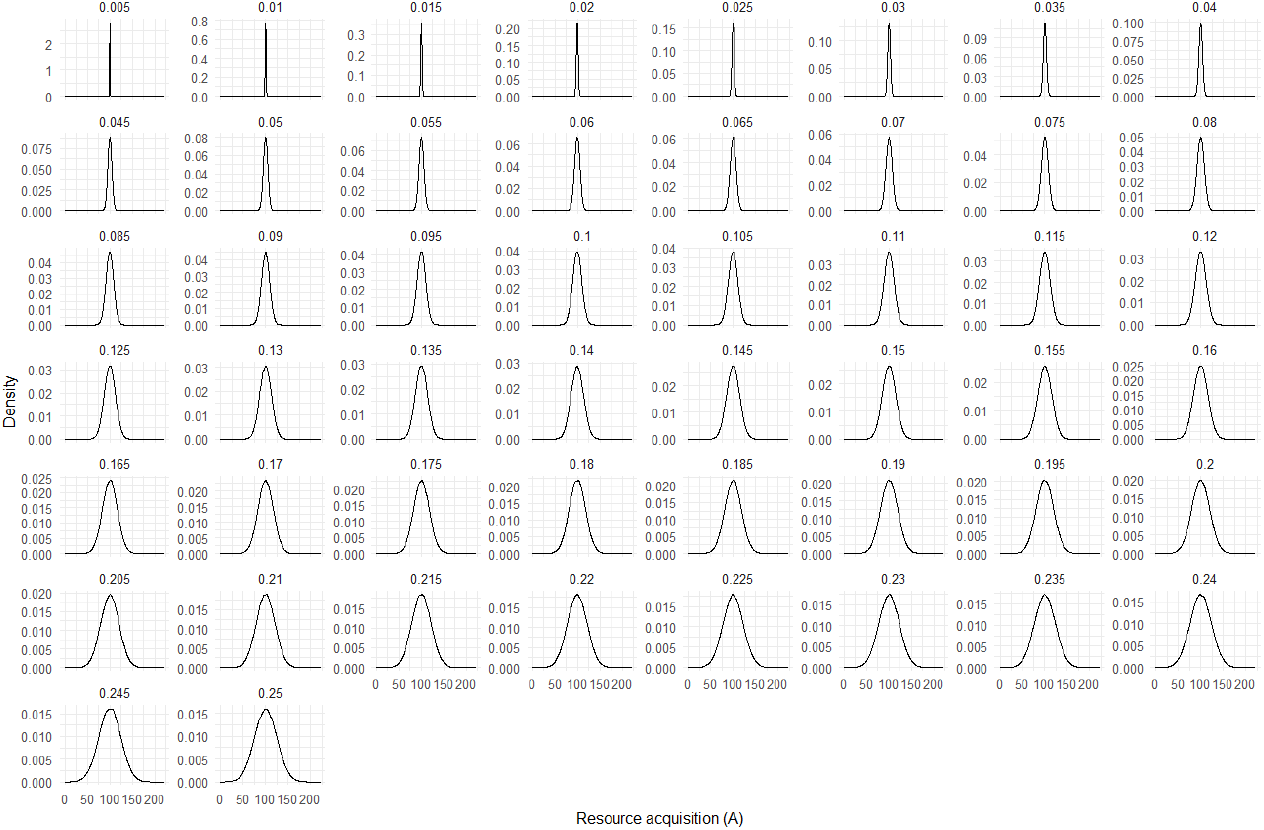
Sample distribution of simulated values for proportion of resources acquired (*A*_*i*_) in model 1. Each facet panel represents different values of CV_*A*_.

**Figure S6:**
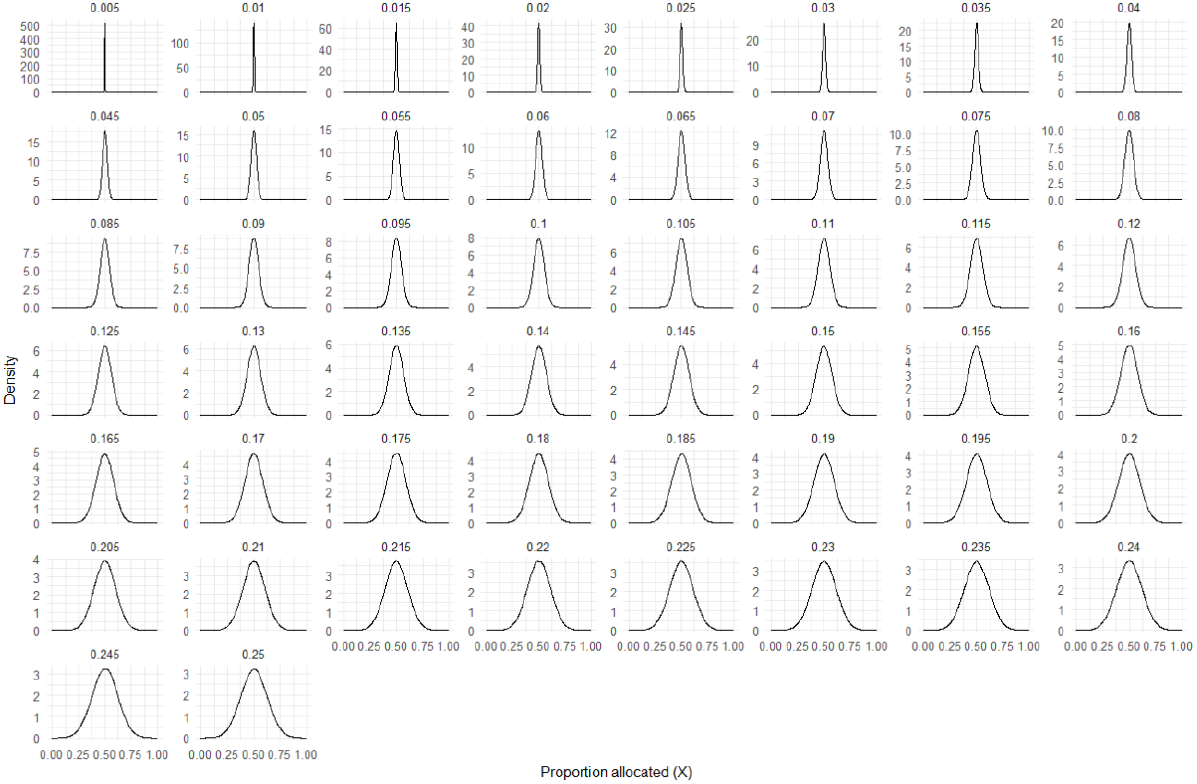
Sample distribution of simulated values for proportion of resources allocated (*X*_*i*_) in model 1. Each facet panel represents different values of CV_*X*_.

**Figure S7:**
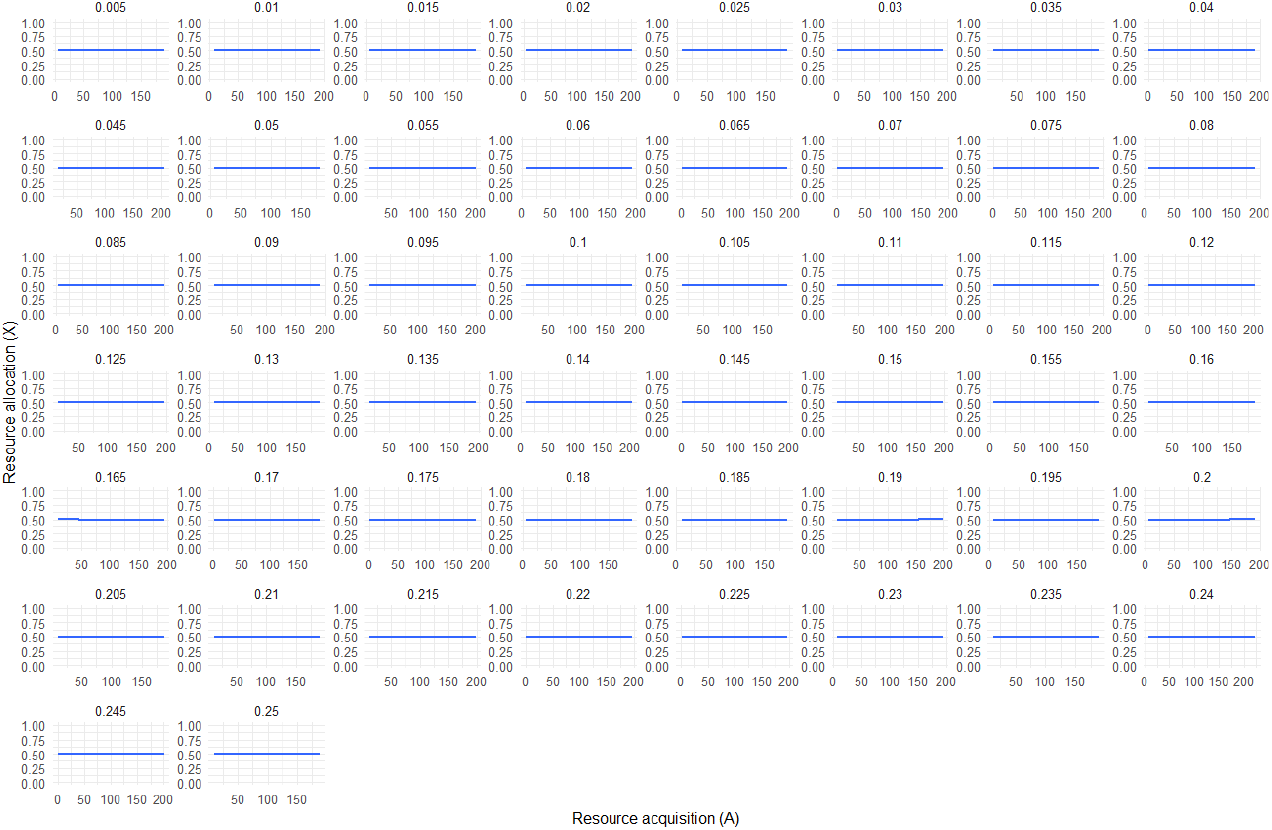
Diagnostic for model 1 showing that there was no covariance between sampled values of *A*_*i*_ and *X*_*i*_. Each facet panel shows a different simulated value of CV_*X*_.

**Figure S8:**
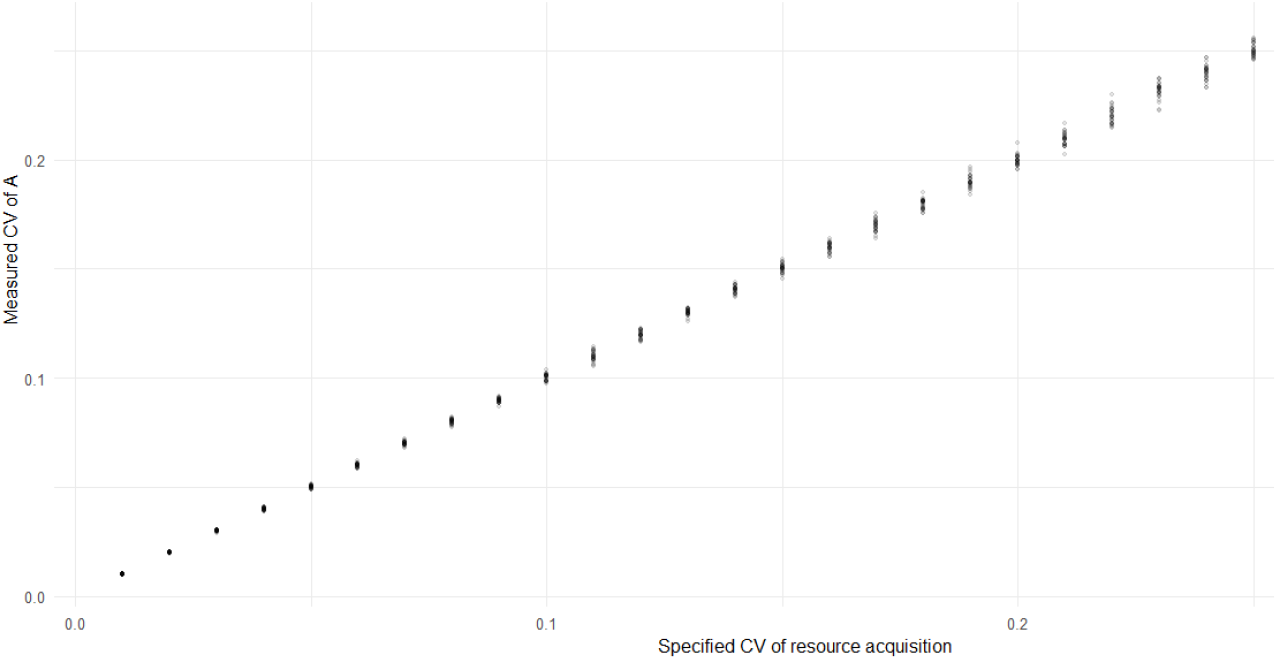
Diagnostic for model 1 showing that the CV_*A*_ values specified corresponded to the simulated CV_*A*_ values. Each dot represents a unique combination of CV_*A*_ and CV_*X*_ simulated using 2500 individuals.

**Figure S9:**
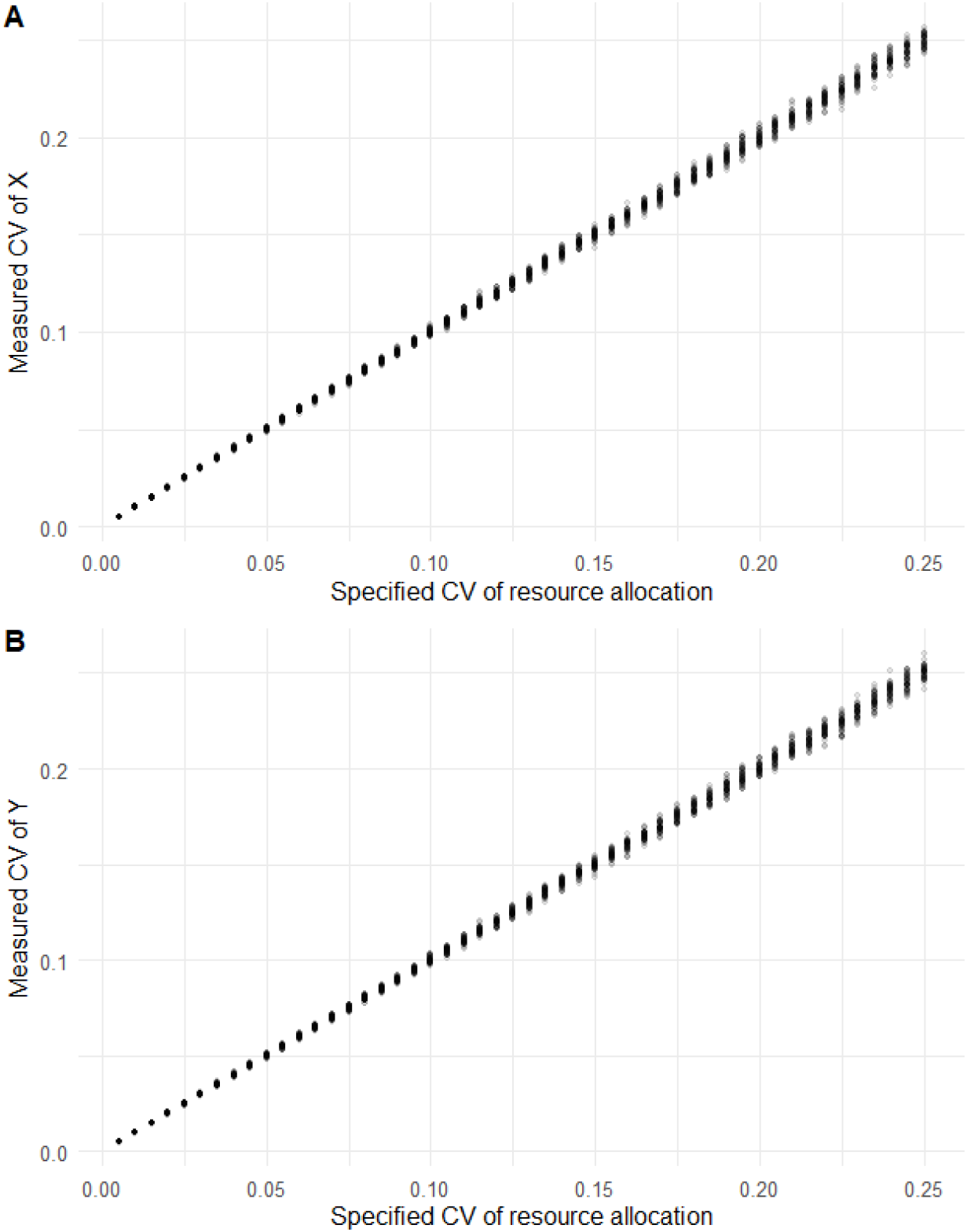
**(A)** Diagnostic for model 1 showing that the specified CV_*X*_ values corresponded to the simulated CV_*X*_ values; and **(B)** to the simulated CV_*Y*_ values.

**Figure S10:**
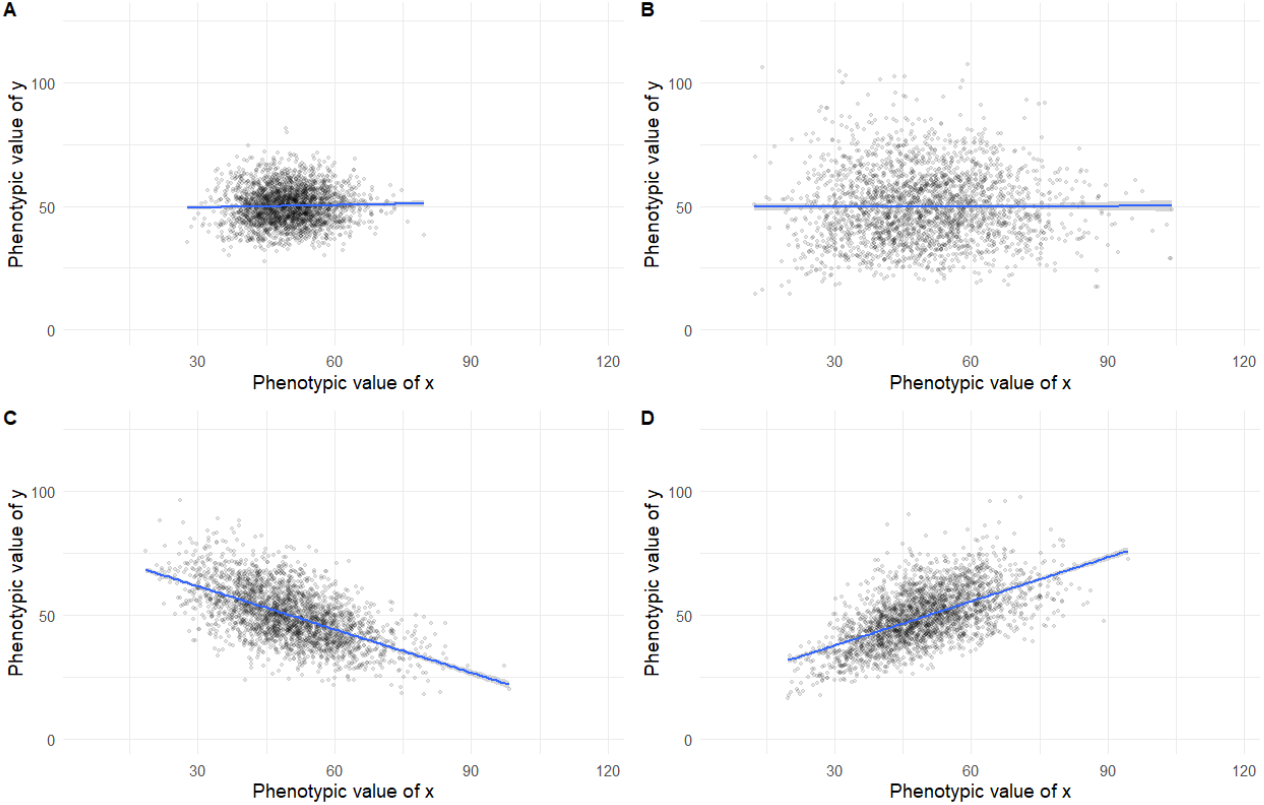
Examples of among-individual phenotypic correlations between traits *x* and *y*, i.e. Cor(*x,y*), in model 1, for different values of CV_*A*_ and CV_*X*_. **(A)** CV_*A*_ = 0.1, CV_*X*_ = 0.1; **(B)** CV_*A*_ = 0.2, CV_*X*_ = 0.2; **(C)** CV_*A*_ = 0.1, CV_*X*_ = 0.2; **(D)** CV_*A*_ = 0.2, CV_*X*_ = 0.1. Each dot represents a single individual, with the phenotypic values of its two traits plotted on the *x* and *y* axes respectively. Each panel consists of 2500 individuals.

**Figure S11:**
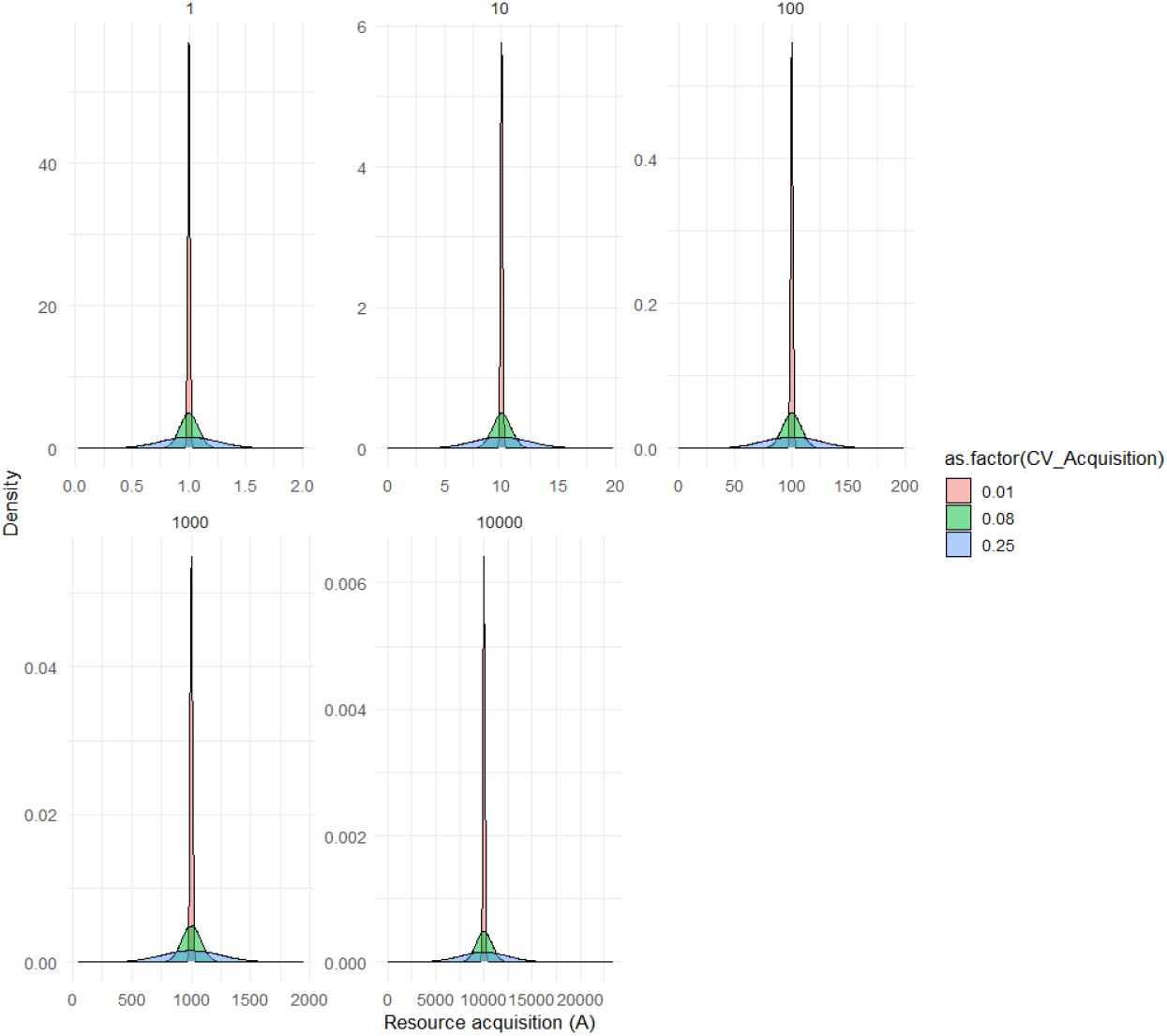
Sample distributions for *A*_*i*_ in model 2A, where different values of μ_*A*_ are simulated (facet panels). Colours of distributions represent some examples of CV_*A*_ values simulated.

**Figure S12:**
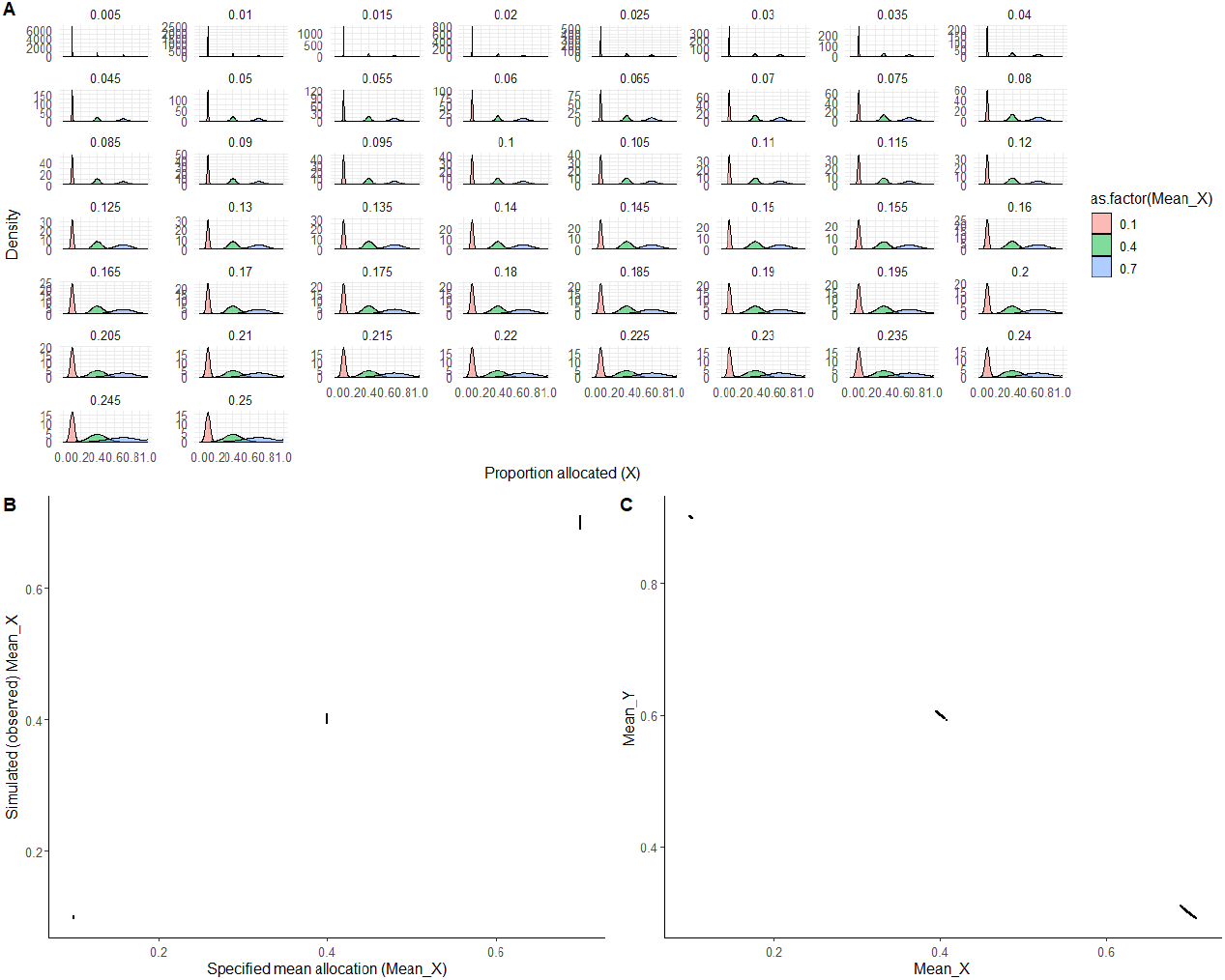
**(A)** Sample distributions for *X*_*i*_ in model 2B, where different values of μ_*X*_ are simulated (colours). Facet panels show different CV_*X*_ values simulated. There was precise agreement between **(B)** specified μ_*X*_ and simulated μ_*X*_ values. **(C)** μ_*X*_ and *μ*_*Y*_ were inversely related, because *Y* = 1-*X*.

**Figure S13:**
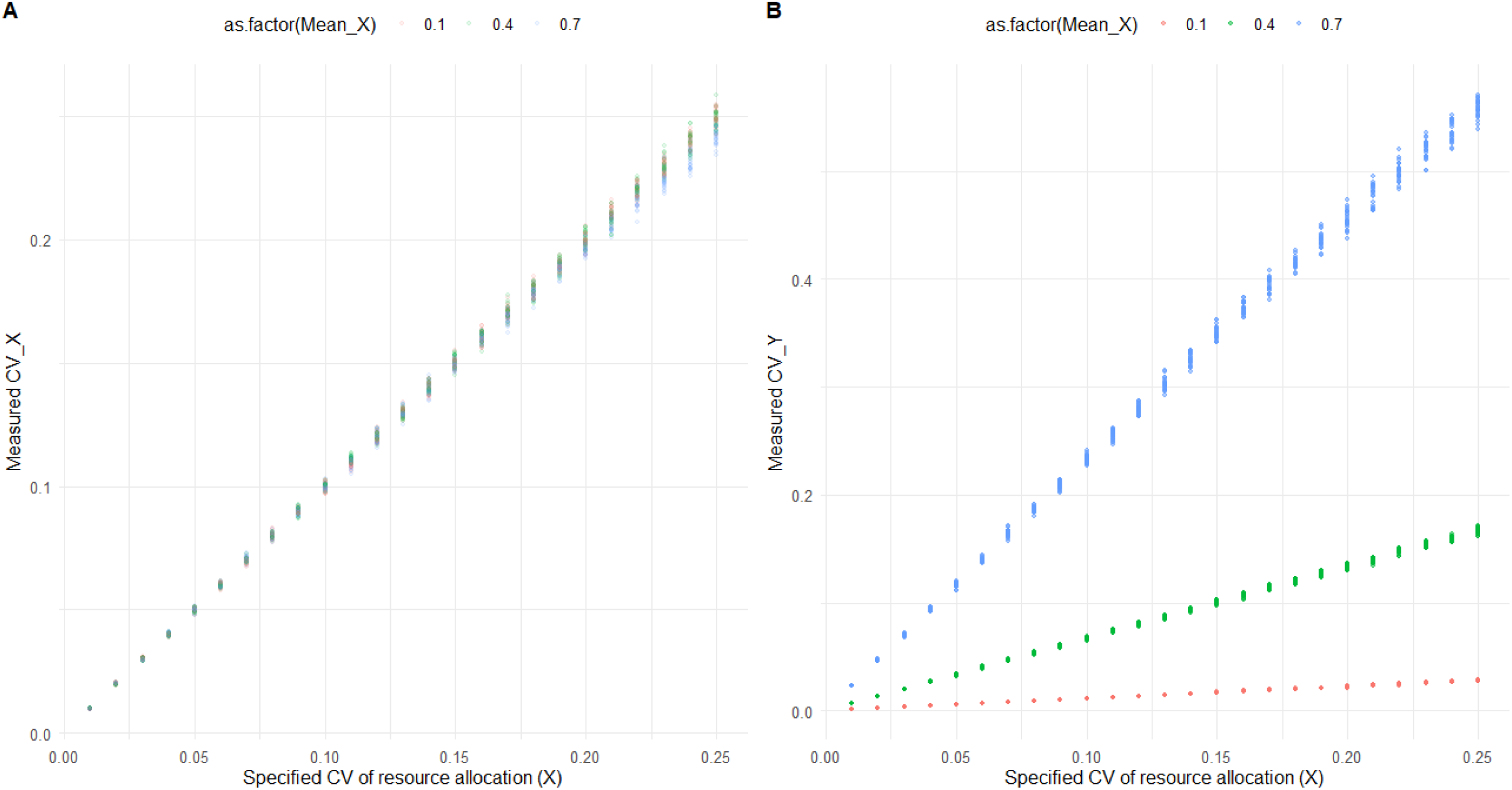
**(A)** Diagnostic for model 2B showing that the CV_*X*_ values specified in the model corresponded to the simulated CV_*X*_ values, irrespective of μ_*X*_. **(B)** Simulated CV_*X*_ values were different from CV_*Y*_ values, and this further depended on μ_*X*_.

**Figure S14:**
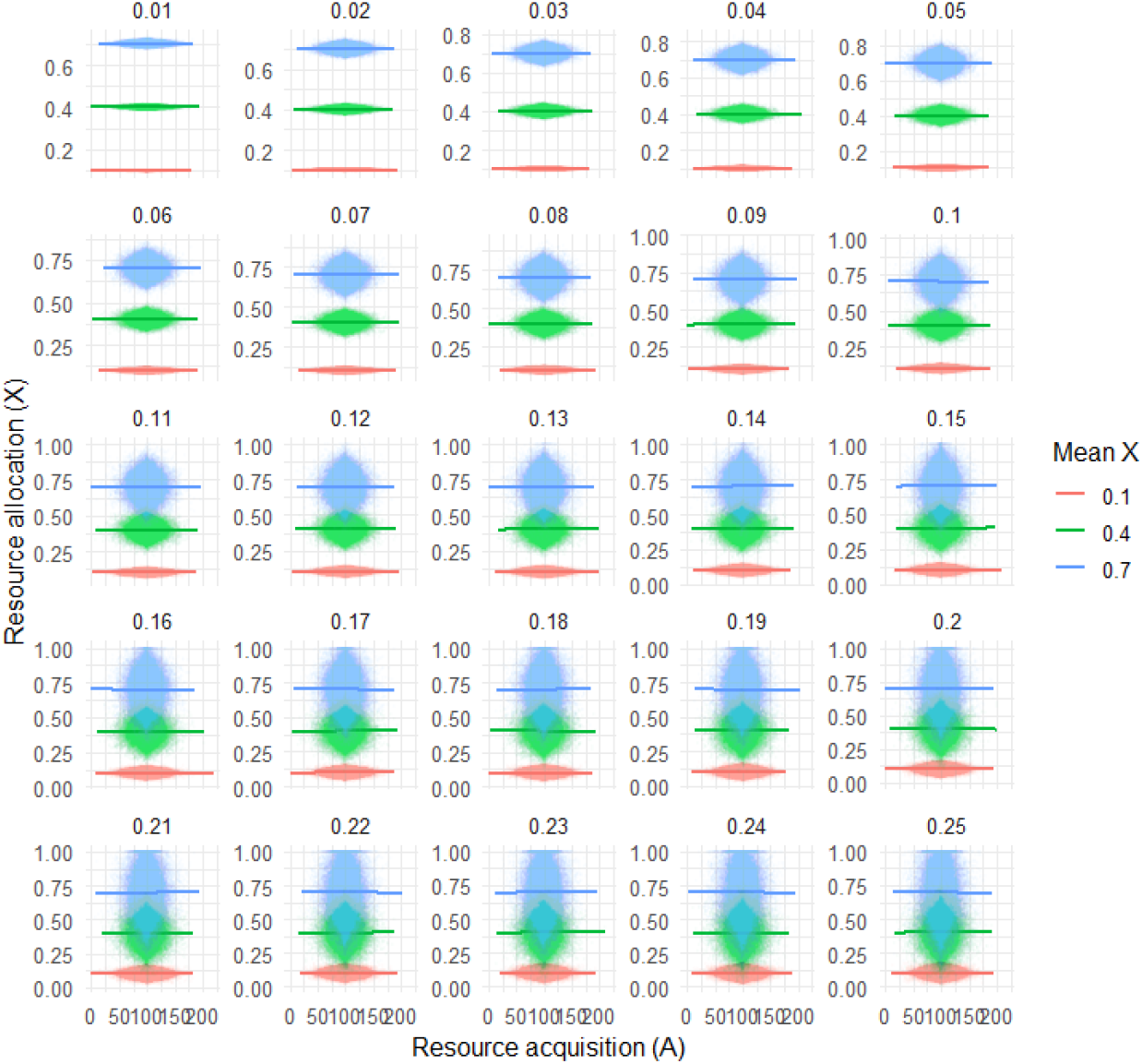
Diagnostic for model 2B showing that there was no covariance between sampled values of *A*_*i*_ and *X*_*i*_. Each facet panel shows a different simulated value of CV_*X*_, each coloured line and corresponding points show a different value for μ_*X*_.

**Figure S15:**
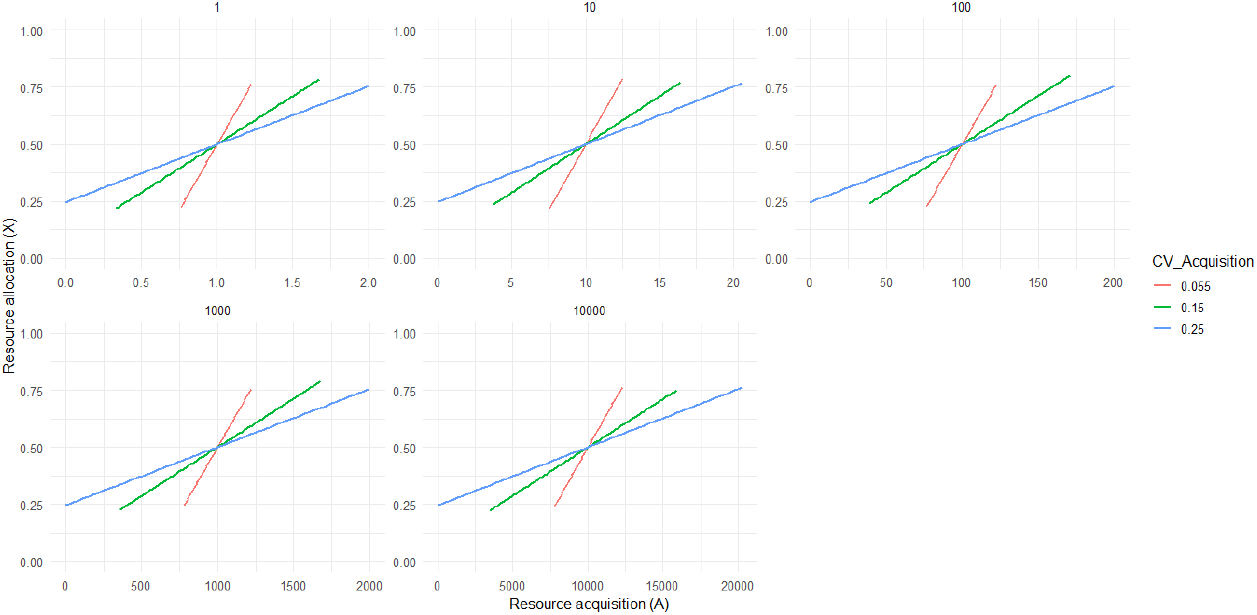
Diagnostic for model 3A showing positive covariance between sampled values of *A*_*i*_ and *X*_*i*_. Each facet panel shows a different simulated value of μ_*A*_, each coloured line shows a different example of CV_*A*_ values simulated.

**Figure S16:**
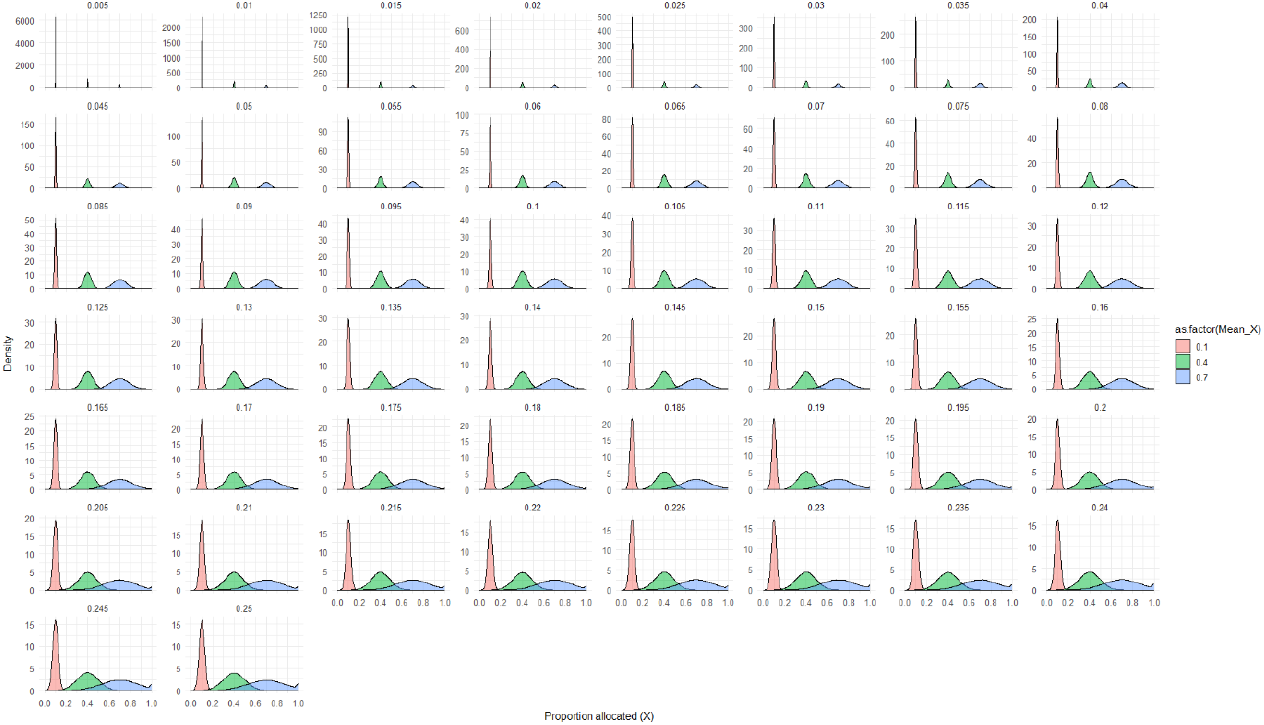
Sample distributions for *X*_*i*_ in model 3B, where different values of μ_*X*_ are simulated (colour). Here, there a positive correlation between acquisition and allocation, i.e. Cor(*A,X*) = 1 was specified. Each facet represents a different CV_*X*_ value.

**Figure S17:**
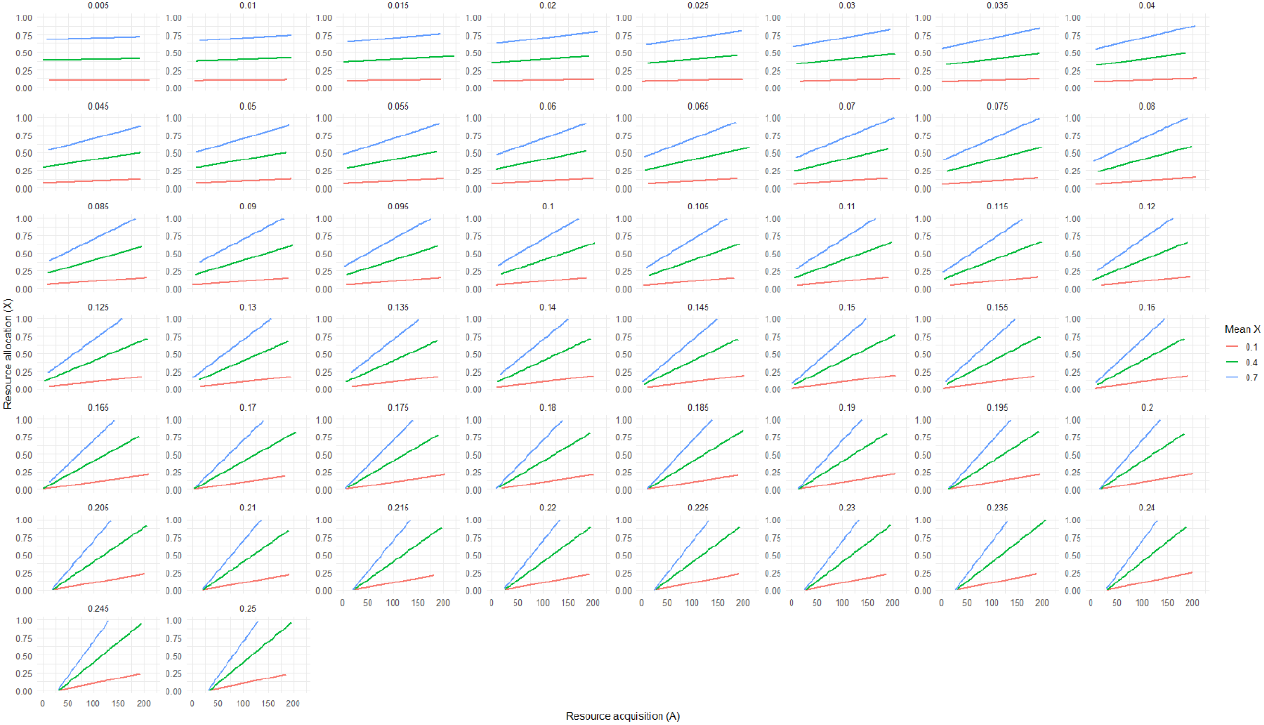
Diagnostics for model 3B, showing positive correlation between resource acquisition (*A*) and proportion allocation (*X*). Facet panels show different CV_*X*_ values, colours show different μ_*X*_ values.

**Figure S18:**
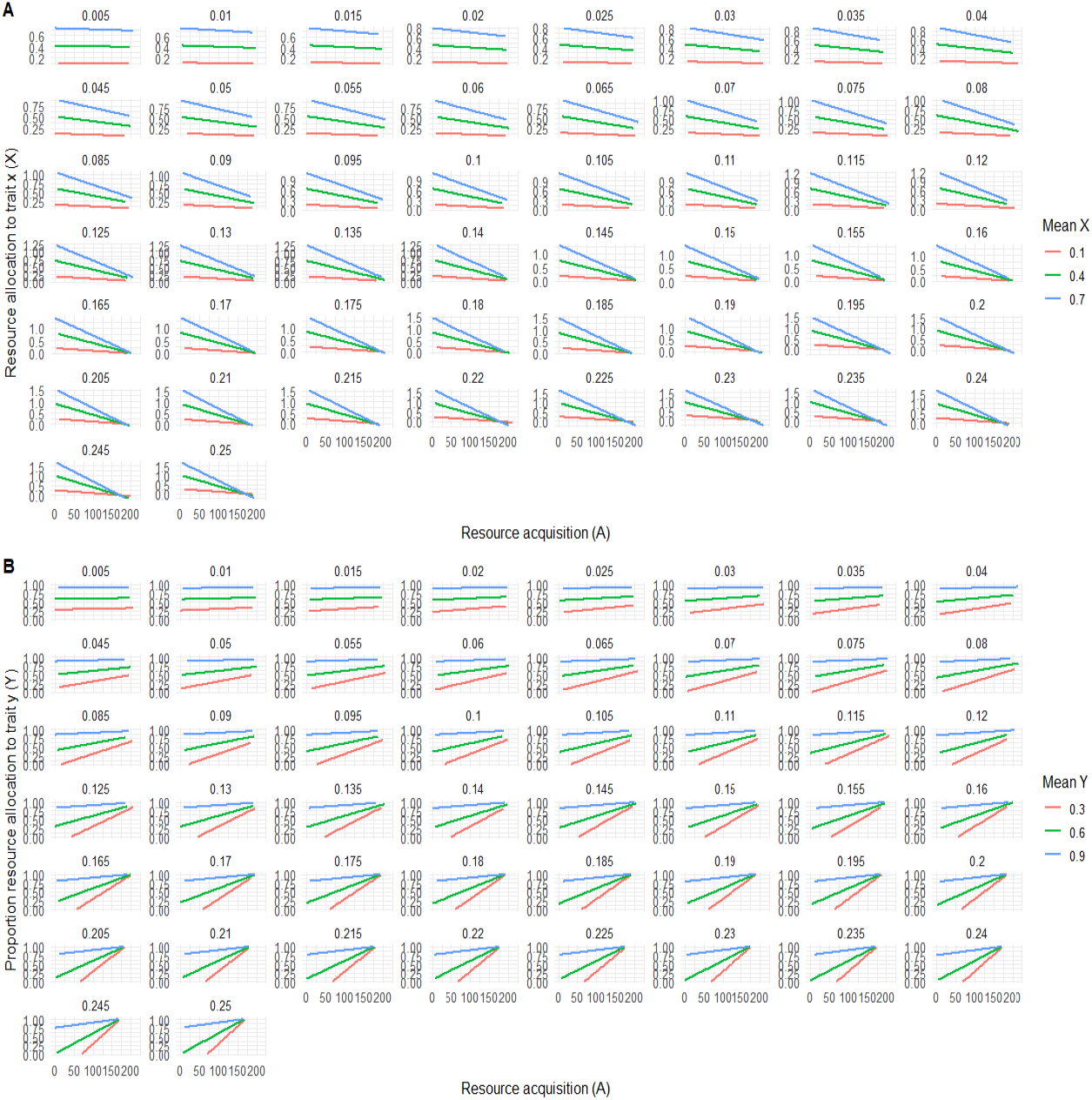
In aim 3B **(A)** negative correlation between *A* and *X* also implies a **(B)** positive correlation between *A* and *Y*. Therefore, correlations between allocation and acquisition have symmetrical meaning, however, requiring swapping of the identity of the trait. In our study, the traits did not have specific biological identity, thus their swapping did not change biological meaning.

**Figure S19:**
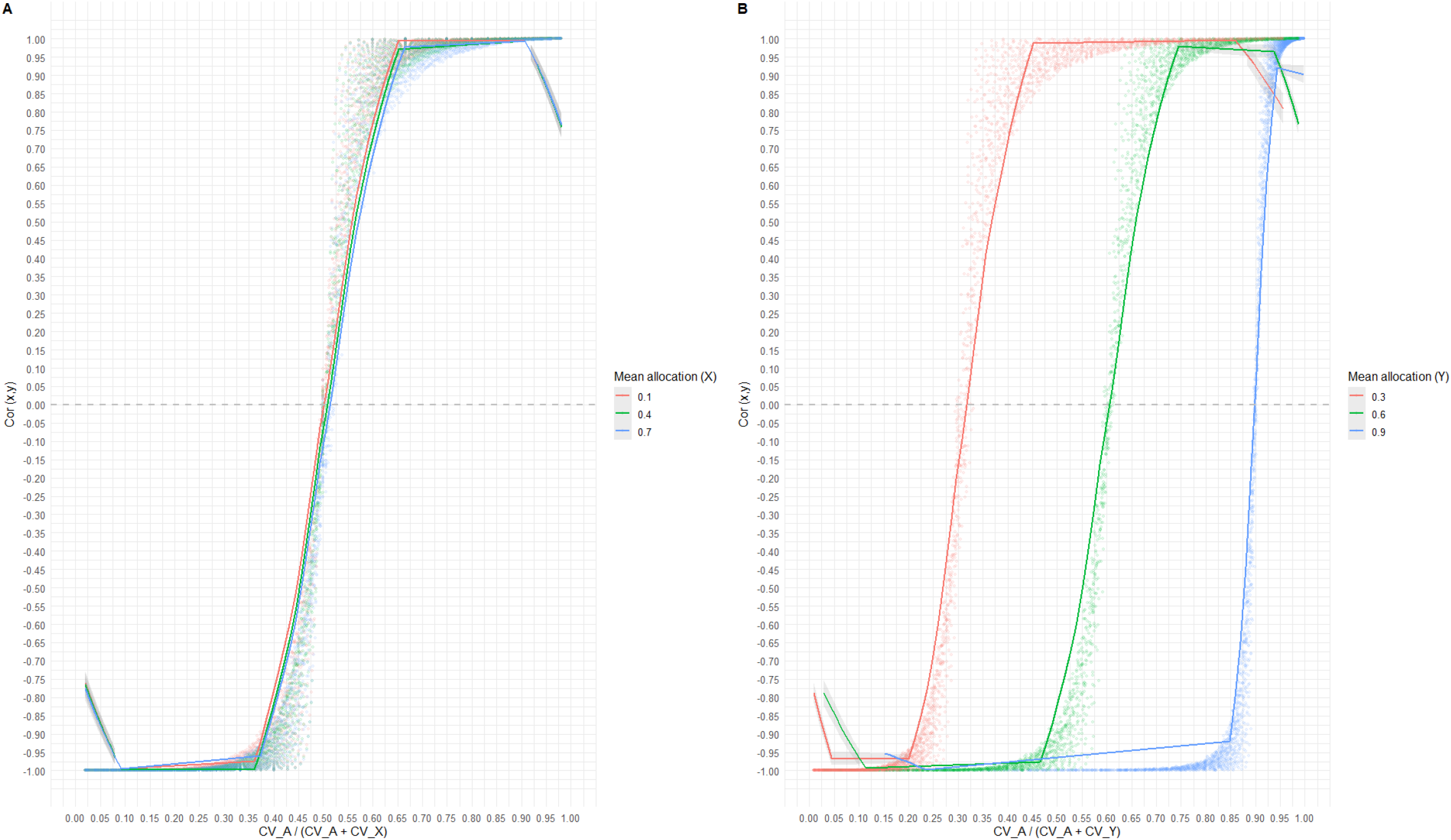
When *A* and *X* covaried negatively, i.e. when *A* and *Y* correlated positively, **(A)** μ_*X*_ did not modulate the influence of CV_*A*_ and CV_*X*_ on Cor(*x,y*). **(B)** However, μ_*Y*_ modulated the influence of CV_*A*_ and CV_*Y*_ on Cor(*x,y*) in a similar way as μ_*X*_ did in model 3B when *A* and *X* correlated positively (see Figure 5). Each dot represents 2500 simulated individuals for a unique combination of CV_*A*_, CV_*X*_, and μ_*X*_ values. Lines show loess smooths.

**Figure S20:**
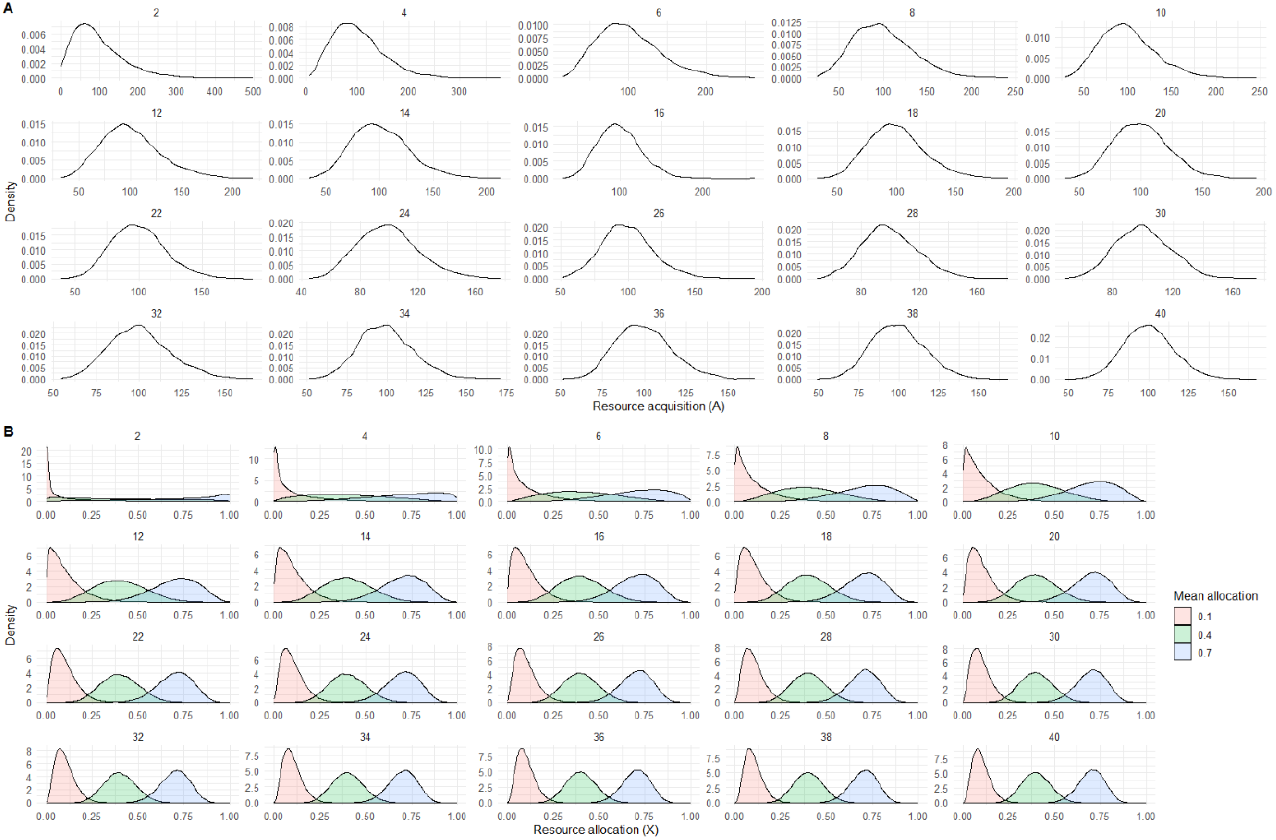
Distribution of **(A)** resource acquisition (*A*_*i*_) and **(B)** proportion allocation (*X*_*i*_), in models where *A*_*i*_ was sampled from a gamma distribution and *X*_*i*_ from a Beta distribution, respectively. CV_*A*_ and CV_*X*_ were retrospectively calculated from the simulated sample distributions, for each unique combination of shape_*A*_, φ, and μ_*X*_. Facet panels show different values of shape_*A*_ in **(A)**, and φ in **(B)**. Colours in **(B)** show different values of mean allocation (μ_*X*_).

**Figure S21:**
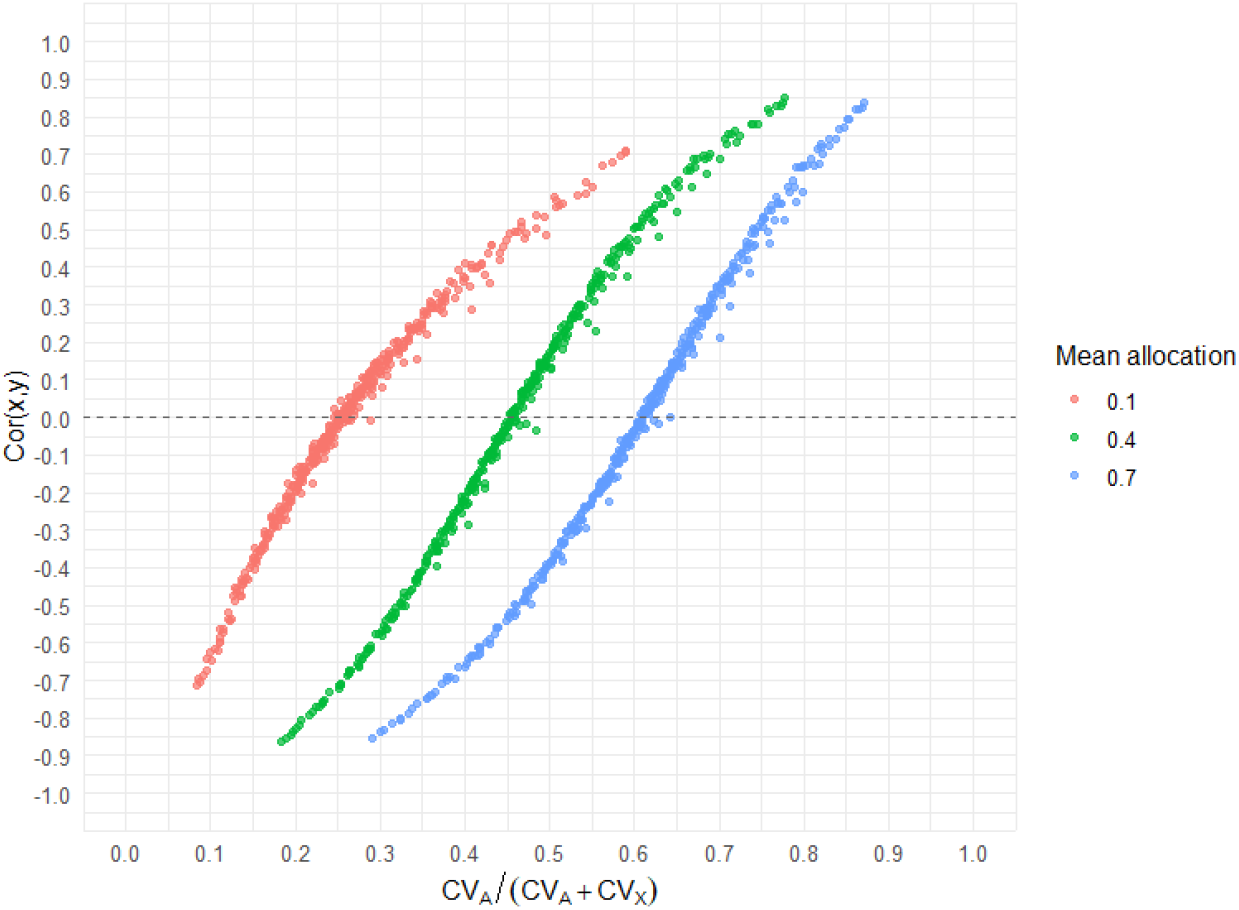
Even in models where *A*_*i*_ was sampled from a Gamma distribution and *X*_*i*_ from a Beta distribution, the phenotypic correlation between traits *x* and *y* was a function of mean allocation (μ_*X*_), and CV_*A*_ and CV_*X*_ (retrospectively calculated from the simulated sample distributions). Each dot, represents 2500 simulated individuals for a unique combination of shape_*A*_, φ, and μ_*X*_.

**Figure S22:**
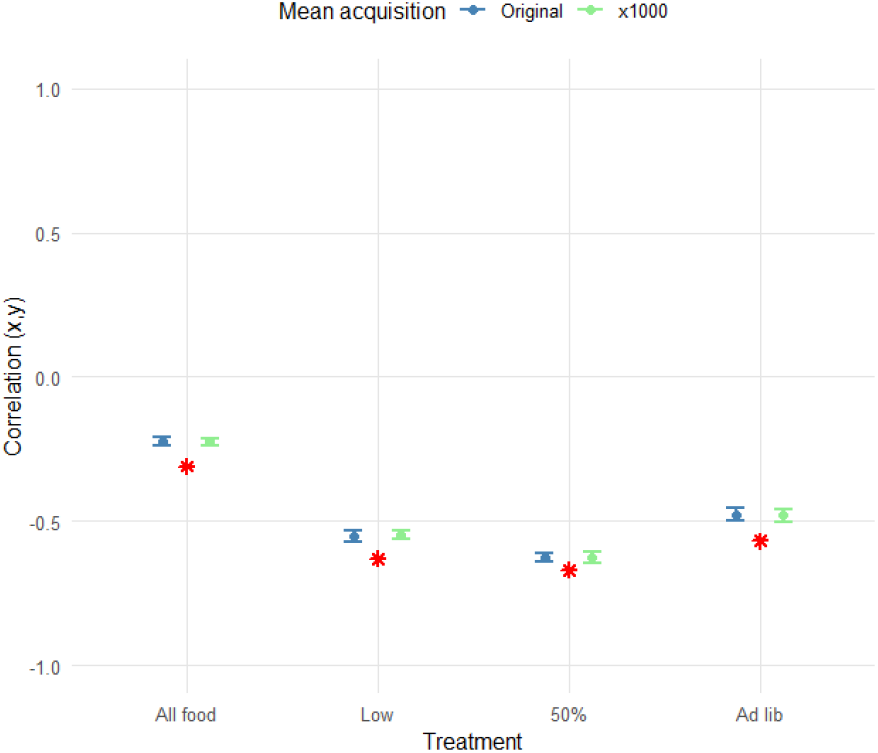
Simulated phenotypic correlation between traits *x* and *y* (dot shows average, error bar shows SD, across 100 replicates) compared against empirically observed phenotypic correlation coefficient (red star) in King et al (2011b), for all four treatments in that paper. In their paper, the traits are flight muscle and ovary mass. Blue values show Cor(*x,y*) when the original reported mean acquisition from King et al was used; green values show Cor(*x,y*) when amplified (x1000) mean acquisition values were used. In each replicate within every simulation, “*N*” number of individuals, as reported in King et al (2011b) were sampled. The axis limits are shown from -1 to +1 to illustrate the entire possible parameter space that Cor(*x,y*) can occupy and the accuracy of our model predictions within it.

**Figure S23:**
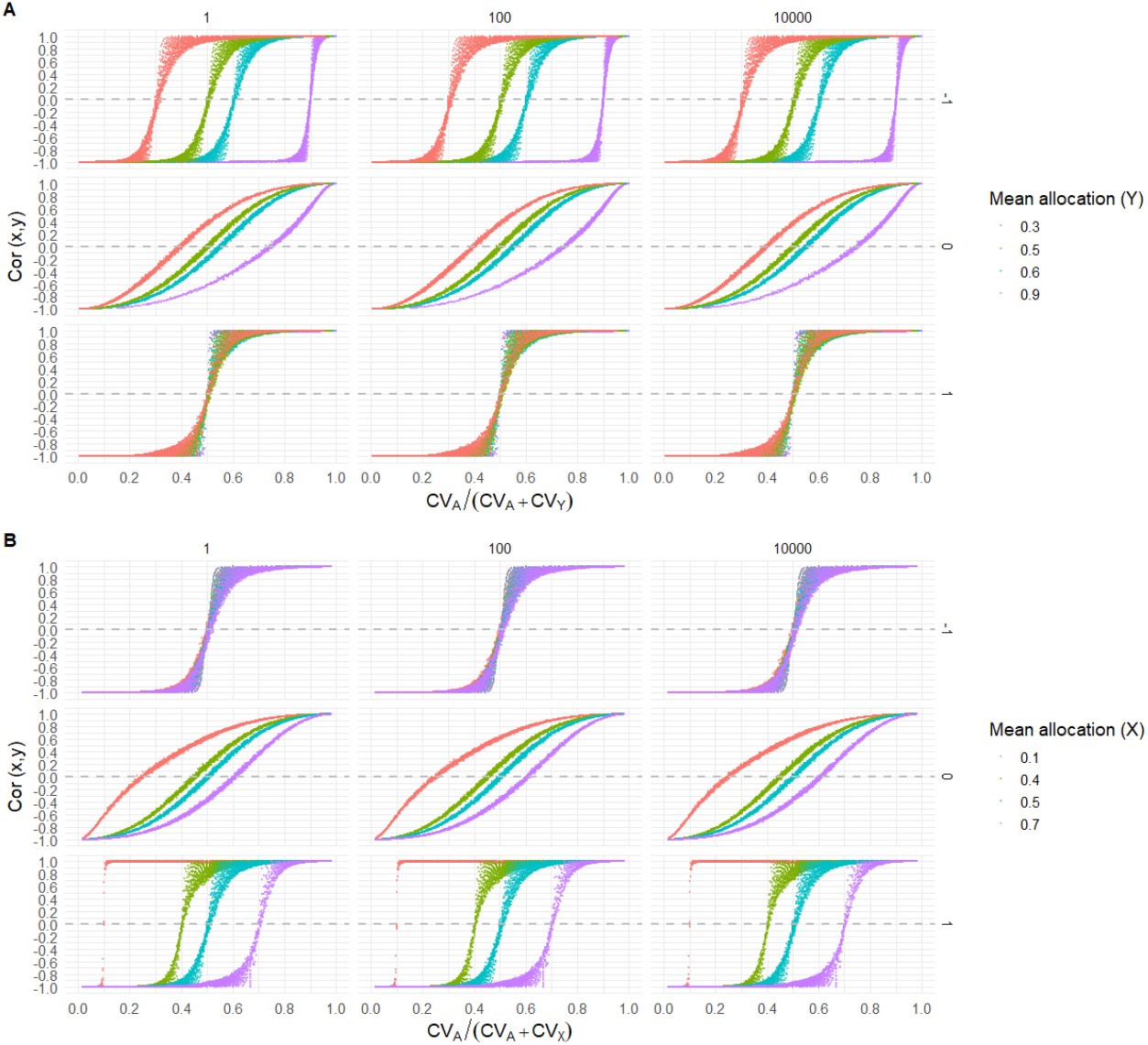
Summary of our results from model 4, where all parameters: μ_*X*_, μ_*A*_, CV_*X*_, CV_*A*_, and Cor(*A,X*), were iterated. **(A)** shows the influence of μ_*Y*_ and CV_*Y*_ while **(B)** shows the influence of μ_*X*_ and CV_*X*_. Column on left represent μ_*A*_ = 1; middle column represents μ_*A*_ = 100, right column represents μ_*A*_ = 10000. Within **(A)** and within **(B)**, rows marked -1, 0, and 1 show Cor(*A,X*) values. Each dot represents a unique combination of μ_*X*_, μ_*A*_, CV_*X*_, CV_*A*_, and Cor(*A,X*) values, and 2500 simulated individuals.

### Supplementary tables

**Table S1:** Summary of parameter values obtained from the “full dataset” section in ‘Table 1’ of King et al (2011b). Variances (σ^2^), standard deviations (σ), coefficients of variation (CV), and means (μ), of acquisition (*A*) and proportion allocation (*X*), as well as the correlation between acquisition and allocation (Cor(*A,X*)), are shown. Traits *x* and *y* in our study represent muscle mass and ovary mass in their study. The values highlighted in grey were used to sample individuals and simulate the phenotypic correlation between traits *x* and *y*. σ and μ were not present in King et al (2011b) and were instead calculated from the presented σ^2^and CV values. *N* is the sample size reported in the study, thus the sample size simulated.

|  | All food | Low | 50% | Ad Libitum |
| --- | --- | --- | --- | --- |
| $N$ | 4148.000 | 1392.000 | 1377.000 | 1379.000 |
| $\sigma^2_A$ | 1.880 | 0.370 | 0.510 | 1.730 |
| $\sigma^2_X$ | 0.068 | 0.067 | 0.060 | 0.049 |
| $\sigma_A$ | 1.371 | 0.608 | 0.714 | 1.315 |
| $\sigma_X$ | 0.261 | 0.259 | 0.245 | 0.221 |
| $CV_A$ | 0.400 | 0.260 | 0.220 | 0.280 |
| $CV_X$ | 0.390 | 0.330 | 0.370 | 0.410 |
| $\mu_A$ | 3.428 | 2.340 | 3.246 | 4.697 |
| $\mu_X$ | 0.669 | 0.784 | 0.662 | 0.540 |
| Amplified $\mu_A$ | 3428 | 2340 | 3246 | 4697 |
| $\text{Cor}(A,X)$ | -0.460 | 0.070 | -0.260 | -0.560 |
| **Empirical $\text{Cor}(x,y)$** | **-0.310** | **-0.630** | **-0.670** | **-0.570** |
| **Simulated $\text{Cor}(x,y)$** | **-0.226** | **-0.551** | **-0.628** | **-0.482** |

**Table S2:** Effect of each parameter on the phenotypic correlation coefficient in a simulation where the parameters: μ_*X*_, μ_*A*_, CV_*X*_, CV_*A*_, and Cor(*A,X*), were all simultaneously varied. Output for the third-best fitting linear model: CV_*A*_/(CV_*A*_ + CV_*X*_) + CV_*A*_/(CV_*A*_ + CV_*Y*_) + μ_*A*_ + μ_*X*_: Cor(*A, X*) illustrates the lack of influence of μ_*A*_.

| Fixed effects | Estimate | SE | t | P |
| --- | --- | --- | --- | --- |
| (Intercept) | -1.872 | 0.010 | -180.516 | **<0.001** |
| $CV_A/(CV_A + CV_X)$ | 0.936 | 0.017 | 53.596 | **<0.001** |
| $CV_A/(CV_A + CV_Y)$ | 2.143 | 0.019 | 111.165 | **<0.001** |
| $\mu_A$ | 0.000 | 0.000 | -0.018 | 0.986 |
| $\mu_X$ | 0.600 | 0.017 | 34.419 | **<0.001** |
| $\text{Cor}(A,X)$ | 0.592 | 0.003 | 181.688 | **<0.001** |
| $\mu_X: \text{Cor}(A,X)$ | -1.186 | 0.007 | -173.558 | **<0.001** |

**Table S3:** Summary of overall results. Parameters that substantially influence the phenotypic correlation between traits *x* and *y*, in each mode marked as ‘Yes’. N/A implies that the parameter was not manipulated in that specific model. Parameter values can be found in Table 1. Also see Figure 7 and S23 for a summary of our results in graphical form.

| Aim<br>(model ID) | $\mu_A$ | $\mu_X$ | $CV_A/(CV_A + CV_X)$ | $Cor(A, X)$ |
| --- | --- | --- | --- | --- |
| 1 | N/A | N/A | **Yes** | N/A |
| 2A | No | N/A | **Yes** | N/A |
| 2B | N/A | **Yes** | **Yes** | N/A |
| 3A | No | N/A | **Yes** | **Yes** |
| 3B | N/A | **Yes** | **Yes** | **Yes** |
| 4 | No | **Yes** | **Yes** | **Yes** |

## Supplementary sections

### Supplementary section S1: Symmetrical relation of *X* and *Y*

In aim 3, we simulated a negative correlation between *A* and *X* (Figure S18A) to show that this was equivalent to a positive correlation between *A* and *Y* (Figure S18B). For this, we specified a negative correlation between the *A* and *X* such that *r* = -1 in equation 9, while keeping the rest of the model the same as model 3B. In this model, while μ_*X*_ did not modulate the influence of CV_*A*_ and CV_*X*_ on Cor(*x,y*), μ_*Y*_ modulated the influence of CV_*A*_ and CV_*Y*_ on Cor(*x,y*) (Figure S19). This result was identical to μ_*X*_ being a modulator when *A* and *X* correlated positively (Figure 5), therefore showing that a positive correlation between acquisition and allocation can be interpreted in the same way as a negative correlation if the identities of the traits are swapped.

### Supplementary section S2: Beta and Gamma distributions

To investigate whether sampling acquisition from Gamma distributions (*A*_*i*_ ∼Γ(shape_*A*_, scale_*A*_), and allocation from Beta distributions (*X*_*i*_ ∼B(α, β)), also produced the relationship: 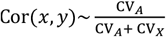, we ran a sensitivity analysis for aim 2B (Figure S20), as proof of concept. In Gamma distributions, the mean of the sample distribution equals scale*shape, where higher shape values imply lower variation. Thus, mean acquisition, μ_*A*_, was fixed at 100 by setting scale_*A*_ = *1*00/shape_*A*_. We iterated shape_*A*_ from 2 to 40 in intervals of 2 to produce various mean-variation combinations. In Beta distribution, φ determines the variation in the data and is calculated as: φ = α + β, where higher φ implies lower variation in the distribution. Thus, for sampling allocation, the Beta distribution was parametrized using μ_*X*_ and concentration (φ) such that: α = φμ_*X*_; β = φ(*1* − μ_*X*_). We iterated different values of φ from 2 to 40 in intervals of 2 to create variation in the data. We also iterated three values for μ_*X*_ as 0.1, 0.3, or 0.7, to produce different mean-variation combinations.

For each unique combination of shape_*A*_, φ, and μ_*X*_, we simulated 2500 individuals. For each of these unique combinations, we retrospectively calculated CV_*A*_ and CV_*X*_ (using Equation 4) from the sampled distributions of *A*_*i*_ and *X*_*i*_ values, as well as calculated Cor(*x,y*). The relationship 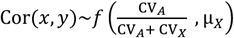 held true even for this sensitivity analyses with more complex *A*_*i*_ and *X*_*i*_ distributions (Figure S21). Simulation code provided on Open science framework: https://osf.io/e89c7/files/osfstorage under the “Beta and gamma” section in the RMD file.

### Supplementary section S3: Empirical verification of model

To test the validity of our simulations and model predictions, we used the data from the classic paper by King et al (2011b) that tests the Y-model, containing data on investment in somatic (muscle) versus gametic (ovaries) tissue, measured as mass. Specifically, we used their data presented in Table 1 containing the phenotypic correlation between the two traits, the CV of allocation and acquisition, and the variance (σ^2^) of allocation of acquisition. We additionally obtained the correlation between acquisition and allocation from page 261 (in text) of their paper. The reported variance and CV were used to calculate the means of allocation and acquisition such that 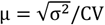. We then used the empirical values of μ_*X*_, μ_*A*_, CV_*X*_, CV_*A*_, and Cor(*A,X*) from their study, to run our simulations.

We conducted four separate simulations, one for each of the food treatments (All food levels, Low, 50%, and Ad Libitum), as presented in their paper. For each simulation, the number of individuals modelled was the same as the sample size “*N*” for that specific diet. In the simulation, we sampled the proportion (*X* and *Y*) and phenotypic values (*x* and *y*) for “*N*” individuals, from normal distributions created using the parameters empirically reported in their study (Table S1). Then, for these “*N*” individuals, we calculated the among-individual phenotypic correlation between the simulated traits *x* and *y*. We ran 100 replicates of each diet-specific simulation and averaged the correlation coefficient across these replicates to then compared this averaged, simulated phenotypic correlation with the ones empirically observed by King et al (2011) (Table S1). The simulated phenotypic correlations were the same sign and similar in magnitude to those observed in the study (Figure S22).

Next, to empirically validate the ‘mean invariance hypothesis’, i.e. mean acquisition (μ_*A*_) not directly impacting phenotypic correlations when other parameters are held constant, we multiplied King et al (2011b)’s reported μ_*A*_ values by 1000, and re-ran each of the four simulations as described above with these amplified μ_*A*_ values. The predicted phenotypic correlation with these amplified μ_*A*_ values produced identical phenotypic correlations to the original simulation values (R^2^ = 0.9896), thus matched the empirically observed phenotypic correlation values by King et al (2011), thereby demonstrating that *μ*_*A*_ has no direct influence on Cor(*x,y*).

To then show that the other parameter (i.e. μ_*X*_, CV_*X*_, CV_*A*_, and Cor(*A,X*)) individually and directly impact the phenotypic correlation, we applied a similar concept. We individually changed the value of one of the other parameters while keeping others the same. As expected, changing values of any parameter other than μ_*A*_ changed the simulated phenotypic correlation.

These analyses are provided on Open science framework: https://osf.io/e89c7/files/osfstorage under the “King et al, 2011” section in the RMD file.

### Supplementary section S4: Analytical validation of model 2B

Under the assumption of no covariance between acquisition and allocation, i.e. Cov (*A, X*) = 0, the phenotypic correlation between traits *x* and *y*, following equation 5 in Van Noordwijk and De Jong (1986), is given by:

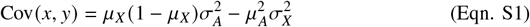

We want to analytically calculate where the zero-crossing point occurs, that is the value of CV_*A*_/(CV_*A*_ + CV_*X*_) where Cor(*x, y*) = 0. Substituting Cor(*x, y*) = 0:

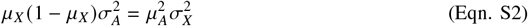

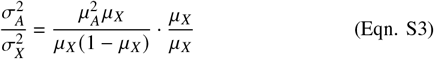

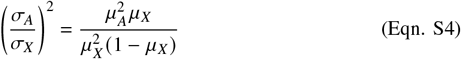

Expressing in terms of coefficients of variation (CV_*A*_ = *σ*_*A*_/*μ*_*A*_, CV_*X*_ = *σ*_*X*_/*μ*_*X*_):

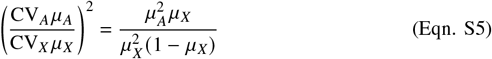

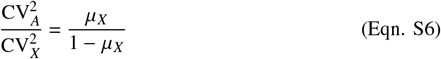

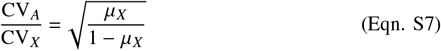

Defining the zero crossing point as 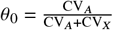. Thus:

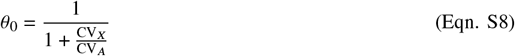

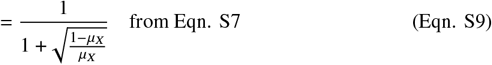

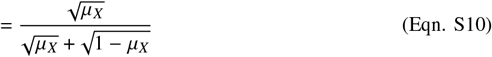

Equation S10 predicts the zero-crossing point for different *μ*_*X*_ values. Comparing analytical predictions to simulation results from model 2B, for where the zero crossing point occurs:

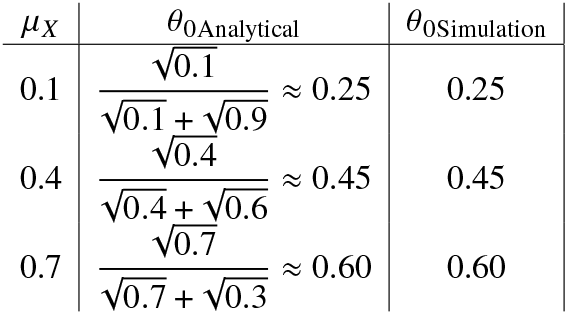

| $\mu_X$ | $\theta_{0\text{Analytical}}$ | $\theta_{0\text{Simulation}}$ |
| --- | --- | --- |
| 0.1 | $\frac{\sqrt{0.1}}{\sqrt{0.1} + \sqrt{0.9}} \approx 0.25$ | 0.25 |
| 0.4 | $\frac{\sqrt{0.4}}{\sqrt{0.4} + \sqrt{0.6}} \approx 0.45$ | 0.45 |
| 0.7 | $\frac{\sqrt{0.7}}{\sqrt{0.7} + \sqrt{0.3}} \approx 0.60$ | 0.60 |

